# Erythropoietin decreases apoptosis and promotes Schwann cell repair and phagocytosis following nerve crush injury in mice

**DOI:** 10.1101/2025.01.22.634402

**Authors:** Prem Kumar Govindappa, Govindaraj Ellur, John P. Hegarty, Akash Gupta, V. G. Rahul, John C. Elfar

## Abstract

After peripheral nerve trauma, insufficient clearance of phagocytic debris significantly hinders nerve regeneration. Without sufficient myelin debris clearance, Schwann cells (SCs) undergo increased apoptosis, impairing functional recovery. There is no treatment for peripheral nerve crush injury (PNCI). Erythropoietin (EPO) is an FDA-approved drug for anemia, which may help in the treatment of PNCI by transdifferentiating resident SCs into repair SCs (rSCs) and enhancing phagocytosis to facilitate the removal of cellular debris. For the first time, we conducted bulk RNA sequencing on mice with calibrated sciatic nerve crush injuries (SNCIs) on days 3, 5, and 7 post-SNCI to uncover transcriptomic changes with and without EPO treatment. We found EPO altered several biological pathways and associated genes, particularly those involved in cell apoptosis, differentiation, proliferation, phagocytosis, myelination, and neurogenesis. We validated the effects of EPO on SNCI on early (days 3/5) and intermediate (day 7) post-SNCI, and found EPO treatment reduced apoptosis (TUNEL), and enhanced SC repair (c-Jun and p75-NTR), proliferation (Ki67), and the phagocytosis of myelin debris by rSCs at crush injury sites. This improvement corresponded with an enhanced sciatic functional index (SFI). We also confirmed these findings *in-vitro*. EPO significantly enhanced SC repair during early de-differentiation, marked by high c-Jun and p75-NTR protein levels, and later re-differentiation with high EGR2 and low c-Jun and p75-NTR levels. These changes occurred under lipopolysaccharide (LPS) stress at 24 and 72h, respectively, compared to LPS treatment alone. Under LPS stress, EPO also significantly increased rSCs proliferation and phagocytosis of myelin or dead SCs. In conclusion, our findings support EPO may enhance the function of rSCs in debris clearance as a basis for its possible use in treating nerve trauma.

## INTRODUCTION

Peripheral nerve crush injury (PNCI) damages myelin and Schwann cells (SCs), leading to long-term morbidity owing to increased inflammation and apoptosis^1,2^. This injury is worsened by the insufficient clearance of myelin and cellular debris at the injury site^3,4^. Nerve regeneration is tightly regulated and begins with resident Schwann cells (SCs) transforming into repair SCs (rSCs)^5,6^. These rSCs then later undergo trans-differentiation into myelin-forming SCs^7^. During this process, rSCs play a crucial role in recruiting macrophages (MΦs)^8^, which initially exist in a pro-inflammatory M1 phase before transitioning to an anti-inflammatory M2 resolution phase^9,10^. The timely transition of both rSCs and MΦs is critical for clearing debris through improved phagocytosis and reduced apoptosis to support injury repair and promote functional recovery.

We demonstrated that erythropoietin (EPO), an FDA-approved treatment for anemia, modulates inflammation and promotes the transition of MΦs from an M1 to an M2 phenotype, enhancing phagocytosis^1^. However, no studies explore the role of EPO in SC transitions, which may be crucial for clearing cellular debris and accelerating recovery. We hypothesized that EPO supports the transition to rSCs to enhance phagocytosis following sciatic nerve crush injury (SNCI), reinforcing our previous finding that EPO improves function after SNCI^1,11–13^. This could only be true if EPO influenced SC activity, and the capacity for phagocytic clearance of broken myelin, in addition to improving the function of M2 MΦs following SNCI.

Few studies investigate novel mechanisms of nerve recovery through functional studies of SCs and MΦs using RNA sequencing^14–24^ in the nerve injury site itself. This relates specifically to phagocytosis, which is critical to prevent nerve recovery problems. Preclinical studies have not resulted into available clinical treatments for nerve trauma, perhaps because of the complexity of cellular responses and post injury transcriptional changes^25,26^. Each cell in the nerve expresses a distinct set of genes that vary based on several factors, including the type and trauma severity^17,24^.

In the present study, using a calibrated mouse SNCI model^27^, we performed transcriptomic evaluation of injured nerve tissue using bulk RNA sequencing. We found that EPO significantly altered several biological pathways and associated genes, especially those related to apoptosis, cell differentiation, phagocytosis, and myelination. We were able to confirm that EPO reduced apoptosis and supported the proliferation of rSCs and phagocytosis of myelin debris in the nerve. This droves myelo-regeneration and promoting functional recovery. EPO also activated M2 MΦs to engulf myelin at the injury site. Our *in-vitro* studies also supported our *in-vivo* observations on nerve tissue regeneration. EPO significantly enhanced early SC de-differentiation (high c-Jun and p75-NTR), proliferation (Ki67), and later re-differentiation (high EGR2 and low c-Jun and p75-NTR) under lipopolysaccharide (LPS) stress conditions. We also demonstrated that EPO significantly increased rSC phagocytosis of both myelin and dead SCs under LPS stress. EPO may be therapeutically useful for nerve trauma as an agent that drives traumatic debris clearance to promote nerve regeneration and function recovery.

## RESULTS

Transcriptomic alterations in injured nerve cells following SNCI were systematically studied using bulk RNA sequencing. Fig. 1 illustrates the experimental processes, including SNCI, EPO dosing, tissue harvesting, RNA extraction, and library preparation for bulk RNA sequencing.

**Fig. 1:**
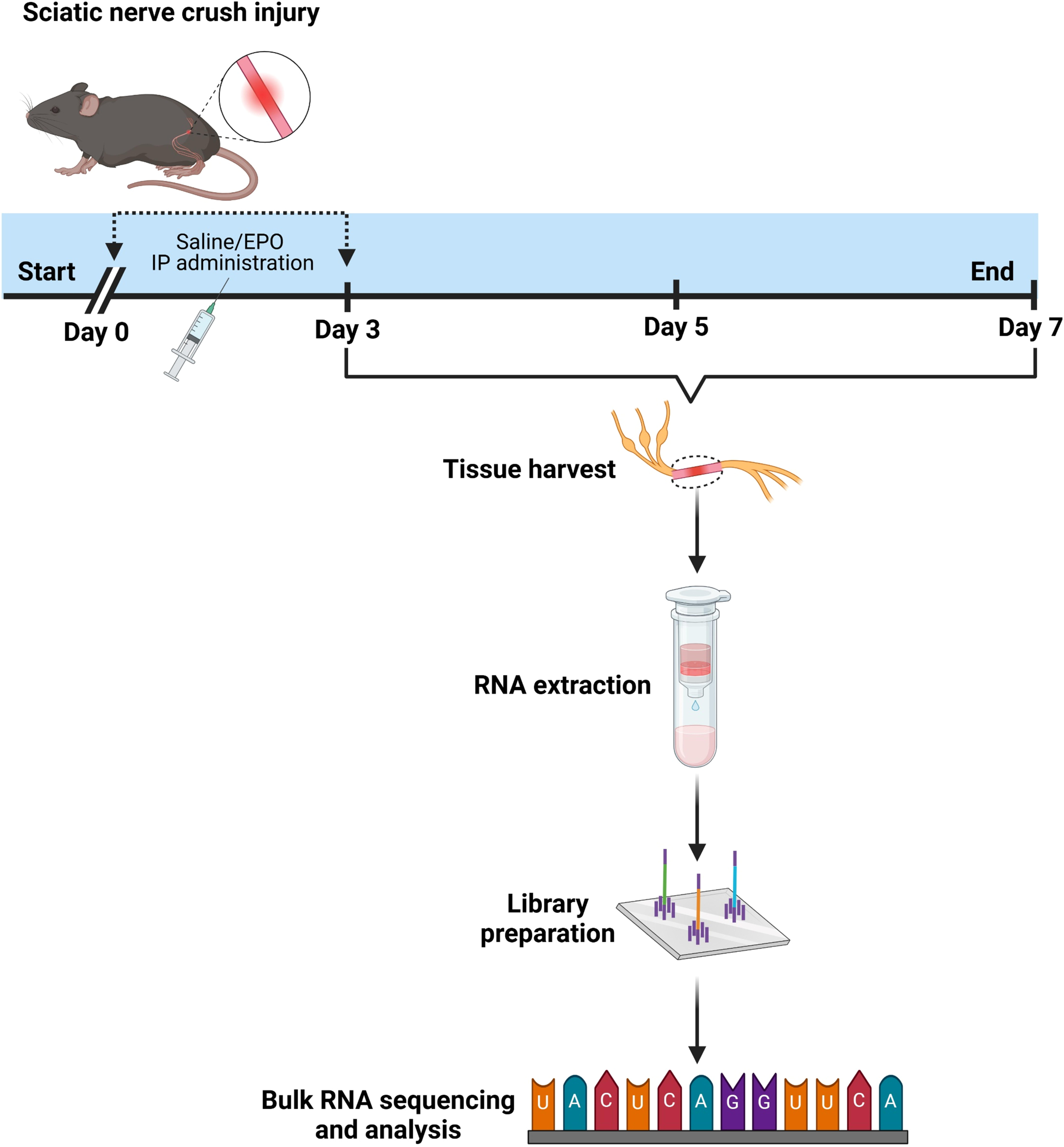
Illustration of the experimental design for bulk RNA sequencing analysis using nerve tissues from saline and EPO treated mice following sciatic nerve crush injury.

### Bulk RNA sequencing showed EPO’s role in enriching genes related to anti-apoptosis, cell differentiation, phagocytosis, and myelination pathways after SNCI

To understand the effects of EPO on cellular transitions and its functional role in regulating various biological pathways, specifically apoptosis, cell differentiation, and phagocytosis following SNCI, we conducted bulk RNA sequencing on post-SNCI days 3, 5, and 7. The principal component analysis (PCA) plot of the transcriptome from nerve tissues treated with EPO showed a distinct cluster compared to the untreated injured group, indicating significant changes in gene expression due to EPO treatment (Figs. 2A, 3A, 4A). Differential gene analysis was conducted using DegSeq to identify differentially expressed genes (DEGs), with a false discovery rate (FDR) of ≤ 0.05, and a fold change greater than 2.0, comparing saline and EPO-treated injured nerves. The results are presented in heat maps (Figs. 2B, 3B, 4B) and volcano plots (Figs. 2C, 3C, 4C).

**Fig. 2:**
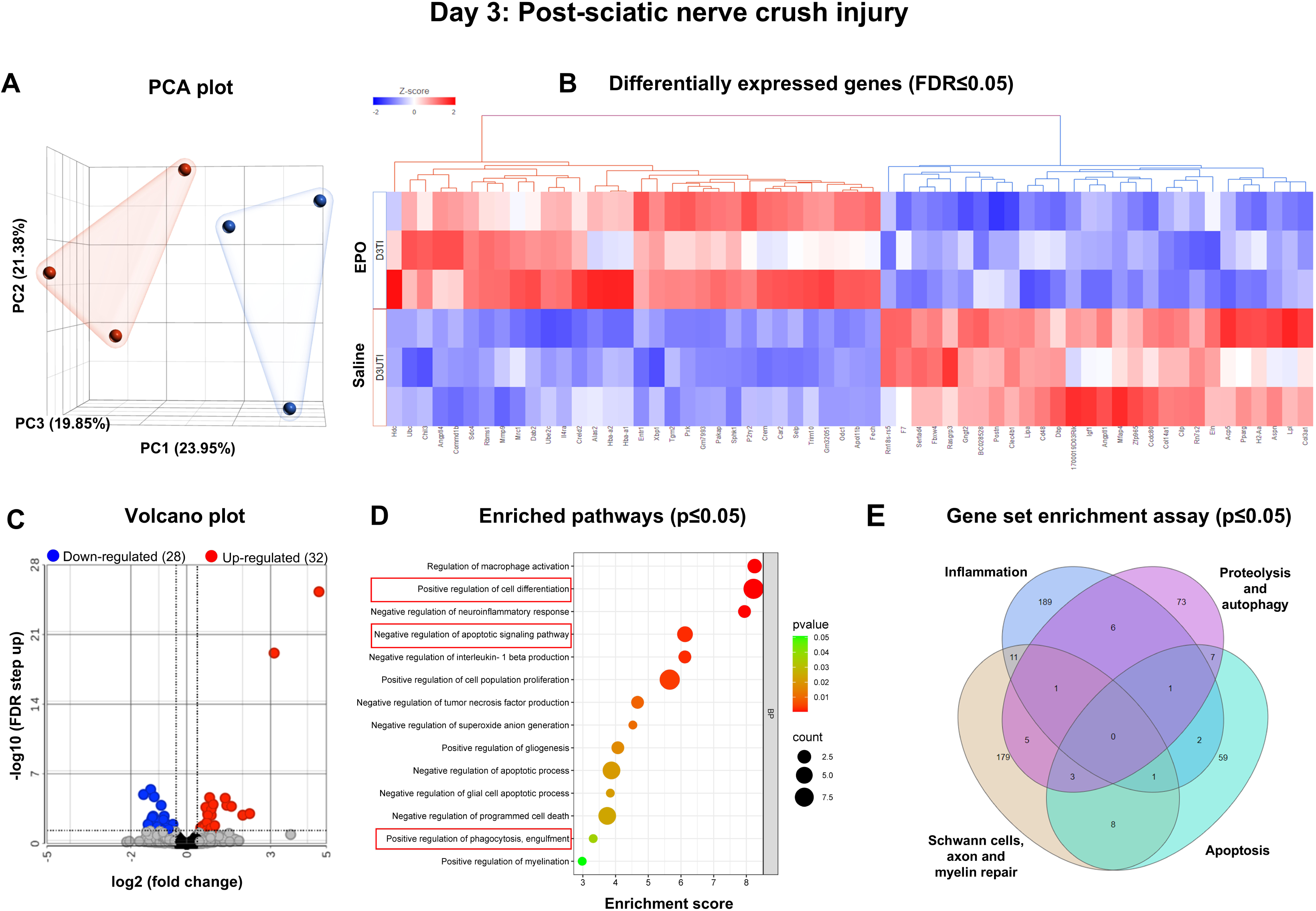
On Day 3, bulk RNA sequencing revealed EPO-enriched genes for biological pathways in nerves following SNCI. **A** Principal component analysis (PCA) plot shows RNA-sequence transcriptomes (saline-red; EPO-blue). **B** Heat map depicts the top upregulated (red) and downregulated (blue) differentially expressed genes (DEGs) (FDR ≤ 0.05). **C** Up and down-regulated DEGs were illustrated in volcano plots to show log 2 (fold change) on the x-axis and significant -log10 (FDR step up) on the y-axis. **D** Enriched pathways from the DEGs are represented on the y-axis with their associated gene numbers, while the x-axis displays the enrichment score for each pathway. **E** The Venn diagram illustrates the number of significantly expressed genes (p ≤ 0.05) within pathways identified through a gene set enrichment assay. Saline vs. EPO treatment, n = 3/ group.

**Fig. 3:**
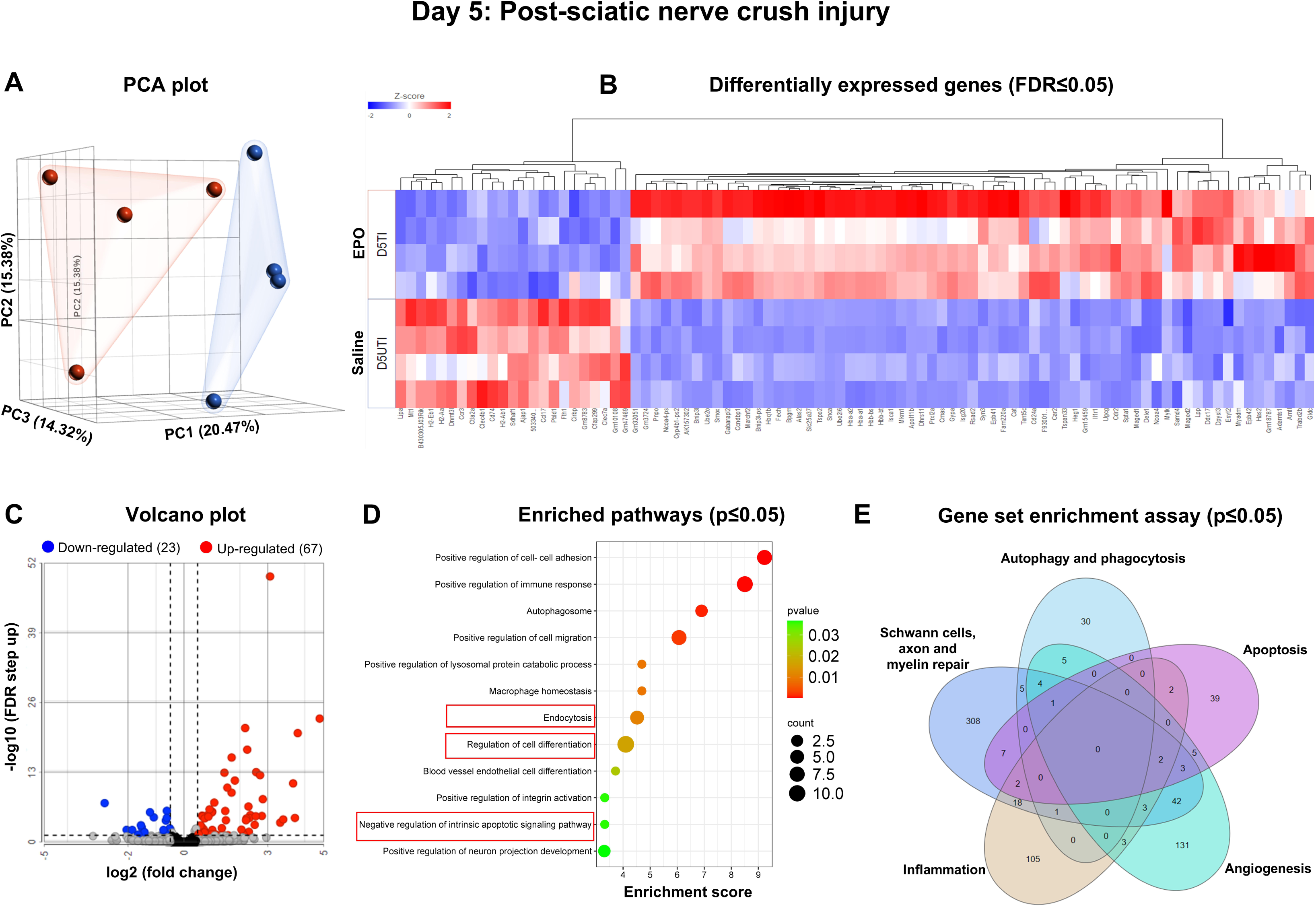
On Day 5, bulk RNA sequencing revealed EPO-enriched genes for biological pathways in nerves following SNCI. **A** Principal component analysis (PCA) plot shows RNA-sequence transcriptomes (saline-red; EPO-blue). **B** Heat map depicts the top upregulated (red) and downregulated (blue) differentially expressed genes (DEGs) (FDR ≤ 0.05). **C** Up and down-regulated DEGs were illustrated in volcano plots to show log 2 (fold change) on the x-axis and significant -log10 (FDR step up) on the y-axis. **D** Enriched pathways from the DEGs are represented on the y-axis with their associated gene numbers, while the x-axis displays the enrichment score for each pathway. **E** The Venn diagram illustrates the number of significantly expressed genes (p ≤ 0.05) within pathways identified through a gene set enrichment assay. Saline vs. EPO treatment, n = 4/ group.

**Fig. 4:**
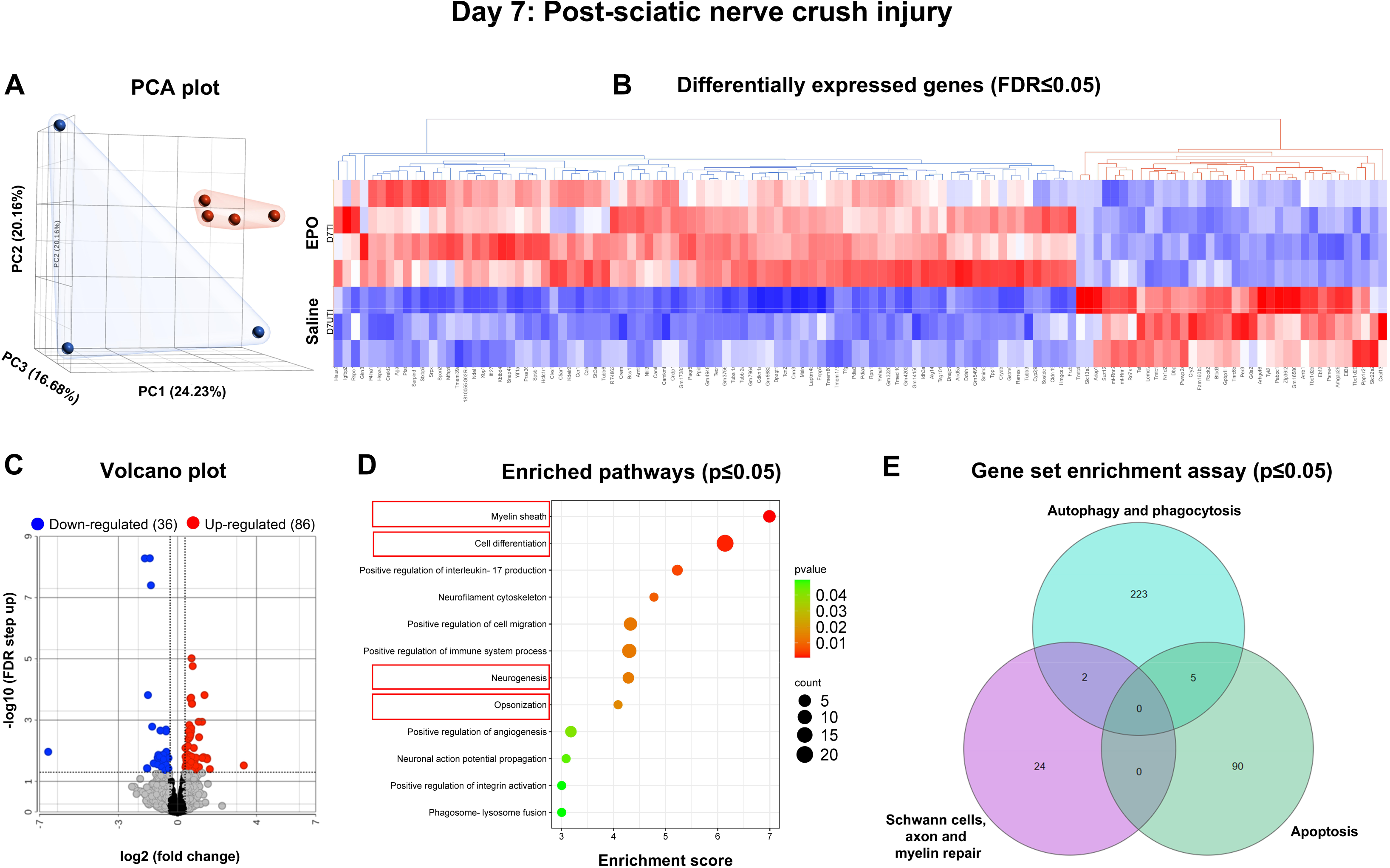
On Day 7, bulk RNA sequencing revealed EPO-enriched genes for biological pathways in nerves following SNCI. **A** Principal component analysis (PCA) plot shows RNA-sequence transcriptomes (saline-red; EPO-blue). **B** Heat map depicts the top upregulated (red) and downregulated (blue) differentially expressed genes (DEGs) (FDR ≤ 0.05). **C** Up and down-regulated DEGs were illustrated in volcano plots to show log 2 (fold change) on the x-axis and significant -log10 (FDR step up) on the y-axis. **D** Enriched pathways from the DEGs are represented on the y-axis with their associated gene numbers, while the x-axis displays the enrichment score for each pathway. **E** The Venn diagram illustrates the number of significantly expressed genes (p ≤ 0.05) within pathways identified through a gene set enrichment assay. Saline vs. EPO treatment, n = 3 and 4/ group.

On day 3, we observed an upregulation of 32 genes and a downregulation of 28 genes in the EPO-treated group compared to the saline group (Fig. 2C). Detailed information regarding gene fold changes and statistical analyses is provided in Supplementary Table 1. Fig. 2D shows significantly enriched pathways (p ≤ 0.05) from the DEGs, particularly those related to the negative regulation of apoptosis (e.g., Hdc, Sphk1, Tgm2, Angptl4) and upregulation of cell differentiation (e.g., Dab2, Trim10, P2ry2, Alas2, Fech, Car2, IL4ra) and phagocytosis (e.g., Selp, Mrc1, Chil3, Crem) genes. The gene set enrichment assay analysis indicated that EPO treatment vs. saline significantly altered various biological pathways and the genes relevant to inflammation (e.g., Mrc1, Ccr5, Cd28, Kit), apoptosis (e.g., Nrg1, Hyou1, Nr4a2, Bcl2l1, Dab2), proteolysis/ autophagy (e.g., Nod1, Gclc, Tmem39a, Snx18, Psen1), and the repair of Schwann cells, axons, and myelin (e.g., Cxcl5, Car2, Gprc5a, Nrg1, Crem, Ngf, Bcl3) (Fig. 2E, Supplementary Fig. 1A-D, Supplementary Table 2).

On day 5, the EPO-treated group showed upregulation of 67 genes and a downregulation of 22 genes compared to the saline group (Fig. 3C). Detailed information on gene fold changes and statistical analyses are provided in Supplementary Table 3. Fig. 3D highlights important enriched pathways (p ≤ 0.05) originating from the DEGs, particularly those related to the negative regulation of apoptosis (e.g., Tspo2, Mt1, Dele1, Prx12a, Spta1), and the positive regulation of cell differentiation (e.g., Car2, Ube2l6, Maged1, Fam220a, Cdr2, Samd4, Alas2, Epb41), and endocytosis/ autophagosome formation (e.g., Snca, Dpysl3, Gabarapl2, Esyt2, Tspan33, Cmas, Ddx17). In addition, our gene set enrichment assay analysis indicated that EPO treatment significantly altered various pathways, and the genes related to inflammation (e.g., Alas2, Il1b, Cxcr2, Ccl19, Tspan2), apoptosis (e.g., Cxcr2, Ptgs2, Nrg1, Tyro3, Epha4, Casp9, Casp1, Cd44,), autophagy (e.g., Snca, Snx30, Dgkd, Ulk3), phagocytosis (e.g., Ccl19, Pld2, Dab2), angiogenesis (e.g., Il1b, Vegfa, Hey2, Ptgs2, Vcl), and the repair of Schwann cells, axons, and myelin (e.g., Lpar3, Itga8, Fzd8, Nrg1, Tgfa, Wnt2) (Fig. 3E, Supplementary Fig. 2A-E, Supplementary Table 4).

On day 7, we identified 86 upregulated genes, and 36 downregulated genes in the EPO-treated group compared to saline (Fig. 4C). Detailed information regarding the fold changes and statistical analyses is represented in Supplementary Table 5. Fig. 4D illustrates notable enriched pathways (p ≤ 0.05) from the DEGs, particularly those relevant to opsonization (e.g., Clvs1, Ccr1), cell differentiation (e.g., Gkn3, Gstm6, Tuba1a, Tubb2a, Smim3, Tpp1, Creld2), neurogenesis (e.g., Cndp1, Arntl, Nfil3, Cryab, C5ar1, Tubb3, Laptm4b), and myelin regeneration (e.g., Msln, Xbp1, Cdkn1a, Tsg101, Calr, Hcfc1r1, Ywhah). Gene set enrichment assay analysis confirmed that EPO treatment significantly altered pathways and genes associated with apoptosis (e.g., Nfil3, Prkn, Rad51, Hspa5, Xbp1), phagocytosis (e.g., Spon2, C5ar1, Tmem175, Rab7b, Hspa8), and the repair of Schwann cells, axons, and myelin (e.g., Nefh, Ece2, Nfasc, Bmp4, Btc, Mag, Pmp22) (Fig. 4E, Supplementary Fig. 3A-C, Supplementary Table 6).

In conclusion, analyses of nerve injury sites on days 3, 5, and 7 post-SNCI allowed for a thorough assessment of the transcriptomic changes over time following SNCI with EPO treatment. These findings highlight the significant effects of EPO on various pathophysiological processes, primarily Schwann cell and macrophage differentiation, phagocytosis, anti-apoptosis, and neurogenesis, by regulating several key genes. These data support EPO’s complex role in enhancing tissue regeneration and promoting functional recovery following SNCI.

### EPO attenuated apoptosis following SNCI

The experimental details of the approved SNCI mouse model and *in-vitro* SC culture studies are highlighted in Fig. 5A. Bulk RNA transcriptomic analysis confirmed that EPO decreases apoptosis signaling pathways and affects related gene expression on day 3 (Fig. 2D, Supplementary Tables 1 and 2, Supplementary Fig. 1A), day 5 (Fig. 3D, Supplementary Tables 3 and 4, Supplementary Fig. 2A), and day 7 (Fig. 4D, Supplementary Tables 5 and 6, Supplementary Fig. 3A). To validate the impact of EPO treatment compared to saline, we assessed apoptotic conditions on post-SNCI days 3 and 7 using the DAB-TUNEL staining method. On day 3, EPO vs. saline treatment significantly protected against apoptosis (4.13 ± 0.30 vs. 16.23 ± 0.44; Fig. 5B, C; ****P < 0.0001). By day 7, EPO treatment continued to show protective effects (vs. day 3 EPO) against apoptosis (22.53 ± 5.81 vs. 4.13 ± 0.30; Fig. 5B, C; *P < 0.05), in comparison to a significant increase in apoptosis in saline-treated mice. These results align with our previous immunofluorescence-TUNEL and PI staining^1^. This study confirmed the significant role of EPO in preventing apoptosis, which supports our transcriptomic analysis and may potentially aid in SC repair and nerve regeneration after injury.

**Fig. 5:**
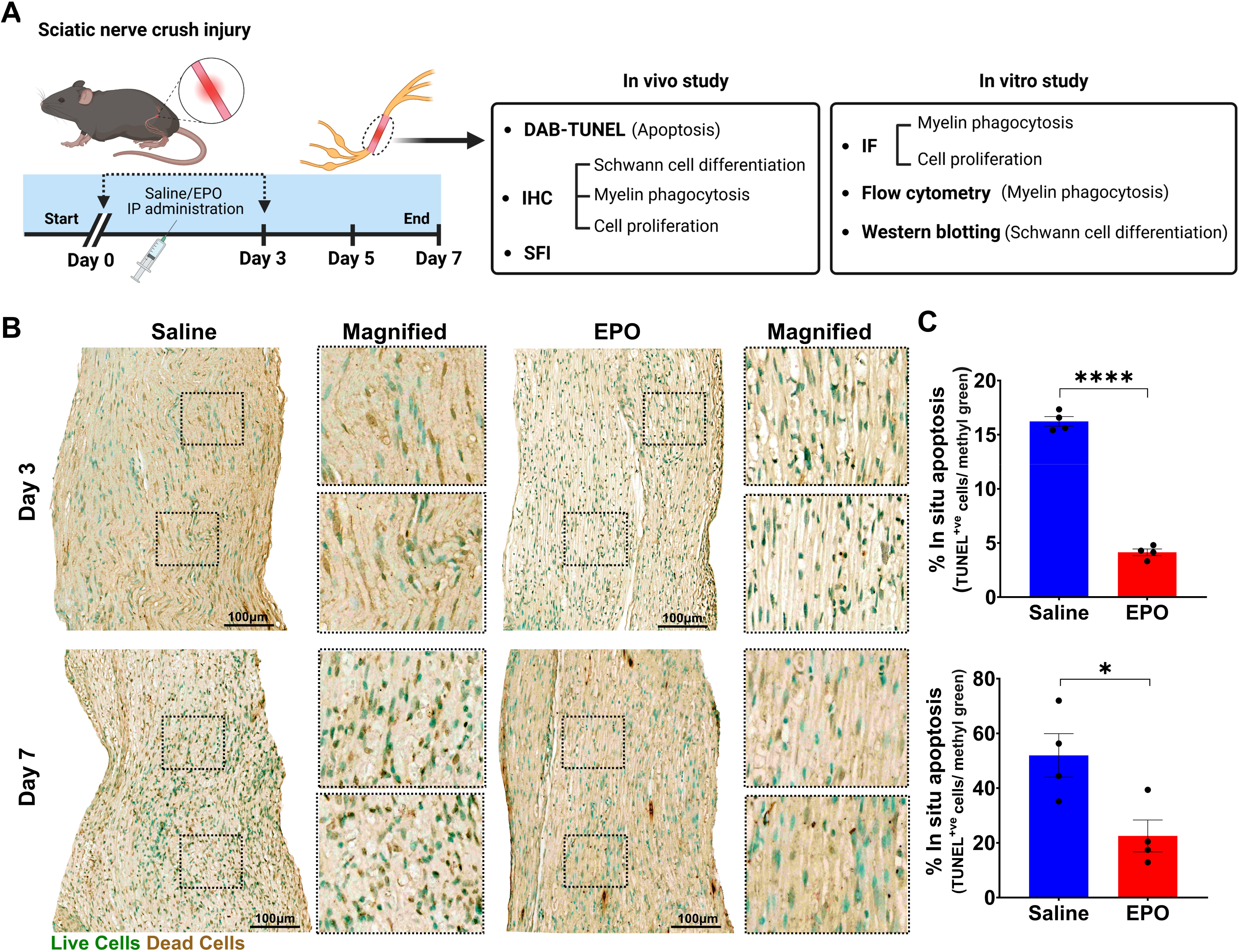
EPO attenuated apoptosis following SNCI. **A** Illustration of the experimental design for *in-vivo* and *in-vitro* cell culture studies. **B, C** Representative images of DAB-TUNEL staining of apoptosis and quantitative results of percent in situ apoptosis (TUNEL positive cells/ methyl green) in saline and EPO treated SNCI tissues on days 3 and 7. n□=□4/ group. Data are represented as mean□±□SEM. The statistical significance is indicated by asterisks (*P□<□0.05 and ****P□<□0.0001 vs. saline group) and compared using two-tailed, unpaired t-tests.

### EPO accelerated Schwann cell repair following SNCI

SNCI obliterates myelin and SCs, but surviving SCs undergo remarkable transcriptional reprogramming to generate repair SCs (rSCs) that express high c-Jun and p75-NTR, and less myelin^28,29^. These rSCs are crucial for phagocytosing cellular debris and later facilitating functional recovery by re-differentiating into myelin SCs^4,30^. Our enriched pathway analysis of DEGs on days 3, 5, and 7 confirmed the significant role of EPO in regulating cell differentiation and myelination pathways and associated genes (Figs. 1D, 2D, 3D, Supplementary Figs. 1C, 2C, 3C, Supplementary Tables 1-6). We aimed to validate EPO’s role in SC repair process following SNCI and under LPS-induced stress conditions, *in-vitro*.

On day 3, IHC analysis revealed that EPO-treated sciatic nerve tissue, compared to the saline group after SNCI, showed a significant increase in the expression of c-Jun (6.95 ± 0.75 vs. 2.65 ± 0.26; Fig. 6A, B; ***P < 0.0002) and p75-NTR (18.88 ± 1.81 vs. 12.00 ± 0.91; Fig. 6A, B; **P < 0.002). However, by day 7, under the same experimental conditions, the expression levels of both c-Jun and p75-NTR had reverted (2.59 ± 0.23 vs. 5.93 ± 0.25 and 6.37 ± 0.60 vs. 14.35 ± 0.73; Fig. 6C, D; ****P < 0.00001). These findings support our hypothesis that EPO promotes early SC dedifferentiation, followed by the transformation of pro-myelin SCs (redifferentiation) with decreased levels of c-Jun and p75-NTR proteins and increased EGR2 expression.

**Fig. 6:**
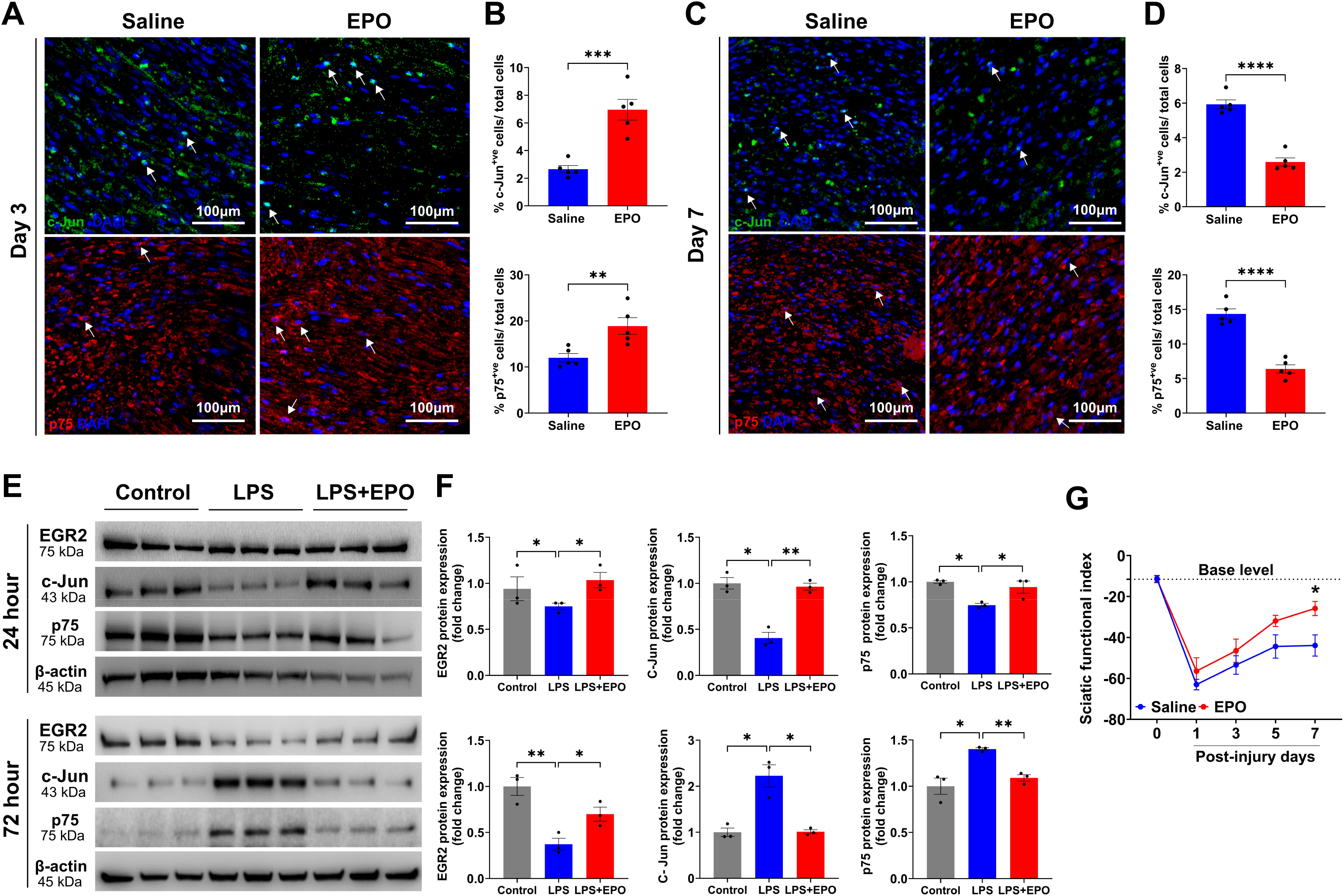
EPO promoted trans-differentiation of Schwann cells and functional recovery following SNCI. Representative IHC images and quantitative results of c-Jun (**A, B**) and p75-NTR (**C, D**) expressions in saline and EPO treated nerve tissues on post-SNCI days 3 and 7. n = 5/ group/ time point. **E, F** Western blotting images and quantitative results of dedifferentiated (c-Jun and p75-NTR) and redifferentiated (EGR2) SCs following 24 and 72h EPO (10IU/ mL) treatment under LPS (500ng/ mL) stress conditions. n = 3/ group. **G** Representation of % sciatic functional index (SFI) untreated vs. EPO treatment. n = 5/ group. Data are represented as mean□±□SEM. The statistical significance is indicated by asterisks (*P□<□0.05, **P < 0.0021, ***P < 0.0002, and ****P□<□0.0001 vs. saline group) and compared using two-tailed, unpaired t-tests or ordinary one-way ANOVA.

Our *in-vitro* study confirmed that EPO treated cultured SCs under LPS stress (vs. LPS alone) conditions significantly increased the expression of c-Jun (0.96 ± 0.03 vs. 0.40 ± 0.06; Fig. 6E, F; **P < 0.0021) and p75-NTR (0.94 ± 0.06 vs. 0.74 ± 0.01; Fig. 6E, F; *P < 0.05) proteins after 24h treatment. After 72h under the LPS stress condition EPO treatment led to early return of expression c-Jun (1.01 ± 0.04 vs. 2.22 ± 0.24; Fig. 6E, F; *P < 0.05) and p75-NTR (1.09 ± 0.03 vs. 1.43 ± 0.01; Fig 6E, F; **P < 0.0021) protein levels as compared to LPS alone treatment, which was no different than what was found in healthy control cells. Also, EPO treatment under LPS stress significantly increased EGR2 protein expression at both 24 and 72h compared to LPS alone (1.03 ± 0.08 vs. 0.74 ± 0.03 and 0.69 ± 0.07 vs. 0.37 ± 0.06; Fig. 6E, F; *P < 0.05). Overall, both *in-vivo* and *in-vitro* data support EPO’s role in promoting SC repair. Our findings suggest that this transformation is essential for the phagocytosis of myelin and cellular debris, which significantly enhances walking function, as measured by the SFI, on day 7 post-SNCI when compared to the untreated saline group (-25.81 ± 3.46 vs. -43.88 ± 5.18; Fig. 6G; *P < 0.05).

### EPO augmented repair Schwann cell and macrophage phagocytosis following SNCI

The clearance of myelin and dead cells through phagocytosis is crucial for reducing inflammation, alleviating cellular stress or apoptosis, and promoting axon regeneration following an SNCI^31,32^. We previously demonstrated that EPO enhances the function of MΦs in clearing fragmented myelin on post-SNCI days 3 and 7 and in cell culture studies under LPS stress conditions^1^. However, we did not identify the peak point of debris clearance with or without EPO treatment in our study. Phagocytosis of debris begins shortly after SNCI and involves both activated rSCs and recruited MΦs. If debris clearance fails, then nerve regeneration is impaired. Based on this understanding, we hypothesized that EPO accelerates the early phagocytosis of cellular debris by rSCs in coordination with the infiltrated MΦ after SNCI. We tested this hypothesis in SNCI and *in-vitro* LPS-induced stress conditions.

On day 3, early after SNCI, IHC results for myelin, using anti-MPZ staining, indicated that damaged myelin at the injury site remained as large myelin fragments. Therefore, rSCs (anti-p75-NTR staining cells) in both the EPO and saline-treated groups showed minimal evidence of phagocytosis (1.24 ± 0.05 vs. 1.16 ± 0.06; Fig. 7A, B; ns). By day 5, there was a noticeable breakdown of large myelin fragments into smaller fragments or debris (Fig. 7A). There was enhanced phagocytosis of rSCs in the EPO-treated group compared to the saline group (51.03 ± 3.19 vs. 29.28 ± 2.34; Fig. 7A, B; ***P < 0.0002). M2 MΦs (anti-CD206 positive staining cells) also exhibited a similar effect with EPO treatment when compared to the saline group (59.53 ± 7.63 vs. 36.44 ± 3.36; Supplementary Fig. 4A, B; *P < 0.05). This suggests an EPO-dependent role for rSCs and M2 MΦs in phagocytosis during the period from day 3 to 5.

**Fig. 7:**
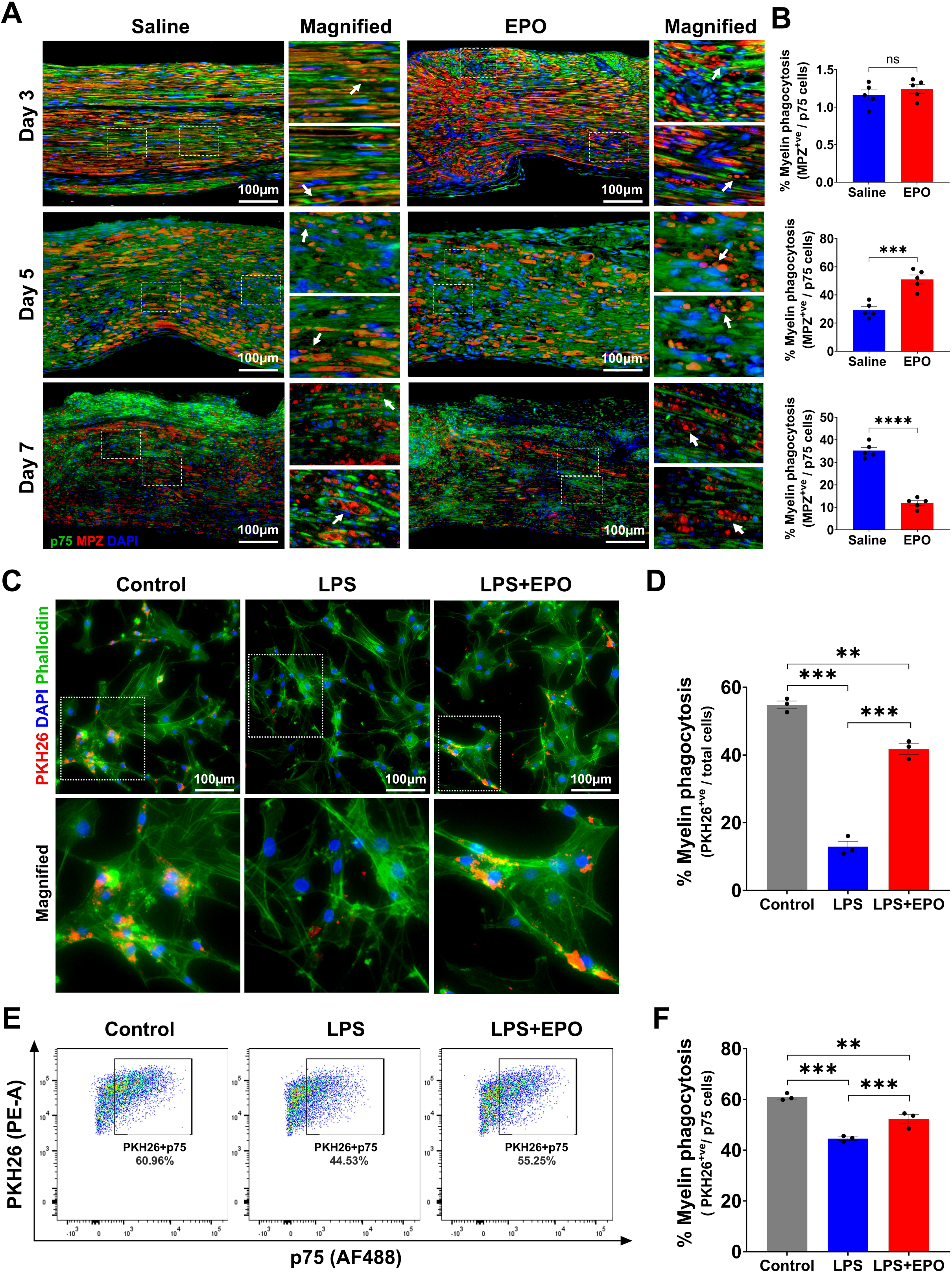
EPO enhanced Schwann cell phagocytosis of cellular debris following SNCI. **A, B** Representative IHC images and quantitative results of repair SCs (anti-p75-NTR staining) phagocytosis of myelin debris (anti-MPZ staining) in saline and EPO treated nerve tissues on post-SNCI days 3, 5, and 7. n = 5/ group. **C, D** Representative IF images and quantitative results of repair SCs (phalloidin staining) phagocytosis of myelin debris (PKH26 staining) following 24h EPO (10IU/ mL) treatment under LPS (500ng/ mL) stress conditions. n = 3/ group. **E, F** Flow cytometry images and quantitative results of repair SCs (p75-NTR positive cells) phagocytosis of myelin debris (PKH26 staining) following 24h EPO (10IU/ mL) treatment under LPS (500ng/ mL) stress conditions. n = 3/ group. Data are represented as mean□±□SEM. The statistical significance is indicated by asterisks (**P < 0.0021, ***P < 0.0002, and ****P□<□0.0001 vs. saline group) and compared using two-tailed, unpaired t-tests or ordinary one-way ANOVA.

On day 7, the EPO-treated group showed a decrease in the percentage of myelin phagocytosis compared to the saline group (11.91 ± 1.03 vs. 35.22 ± 1.46; Fig. 7A, B; ****P < 0.00001). This reduction may be attributed to early clearance of myelin debris (day 5, EPO treatment), suggesting a delayed phase of phagocytosis in the untreated group. We also conducted *in-vitro* experiments to investigate the functional role of EPO in augmenting rSCs phagocytosis using IF and flow cytometry. rSCs treated with EPO under LPS stress conditions displayed a significant increase in myelin phagocytosis compared to those treated with LPS alone (41.75 ± 1.55 vs. 12.96 ± 1.58; Fig. 7C, D; ***P < 0.0002 and 52.20 ± 1.86 vs. 44.53 ± 0.70; Fig. 7E, F; ***P < 0.0002). Healthy control rSCs exhibited characteristic phagocytosis (Fig. 7 C-F). EPO also significantly enhanced the phagocytosis of dead SCs by rSCs under LPS stress conditions compared to LPS alone (54.63 ± 1.20 vs. 31.43 ± 2.72; Supplementary Fig. 5A, B; **P < 0.0021). In our previous publication, we demonstrated similar phagocytic activity using cultured M2 MΦs under LPS stress conditions with the EPO treatment^1^. These data support EPO’s role in augmenting both rSCs and MΦs phagocytosis perhaps through early activation of resident SCs to rSCs and recruitment of MΦs at the injury site, which are crucial for clearing cellular debris. This process may ultimately reduce cell death and promote nerve regeneration and functional recovery.

### EPO increased Schwann cell proliferation following SNCI

Studies have shown that approximately 65 to 80% of the resident cells within the sciatic nerve are SCs, making them the predominant cell type in nerve tissue^33,33,34^. Clearing cellular debris significantly enhances the proliferation of SCs and initiates the remyelination process, which is essential for nerve regeneration following SNCI^1,4^. Our bulk RNA sequencing data on post-SNCI days 3, 5, and 7, revealed a significant role of EPO in promoting cell proliferation pathways and associated genes (Figs. 1D, 2D, 3D). We aimed to confirm EPO’s role in nerve tissue cell proliferation after SNCI on days 3 and 7. IHC staining for Ki67 showed that EPO treatment significantly increased SC proliferation on post-SNCI day 3 (17.59 ± 1.84 vs. 8.38 ± 0.70; Fig. 8A, B; ***P < 0.0002) and day 7 (46.34 ± 2.52 vs. 31.95 ± 3.20; Fig. 8A, B; *P < 0.05) compared to saline treatment.

**Fig. 8:**
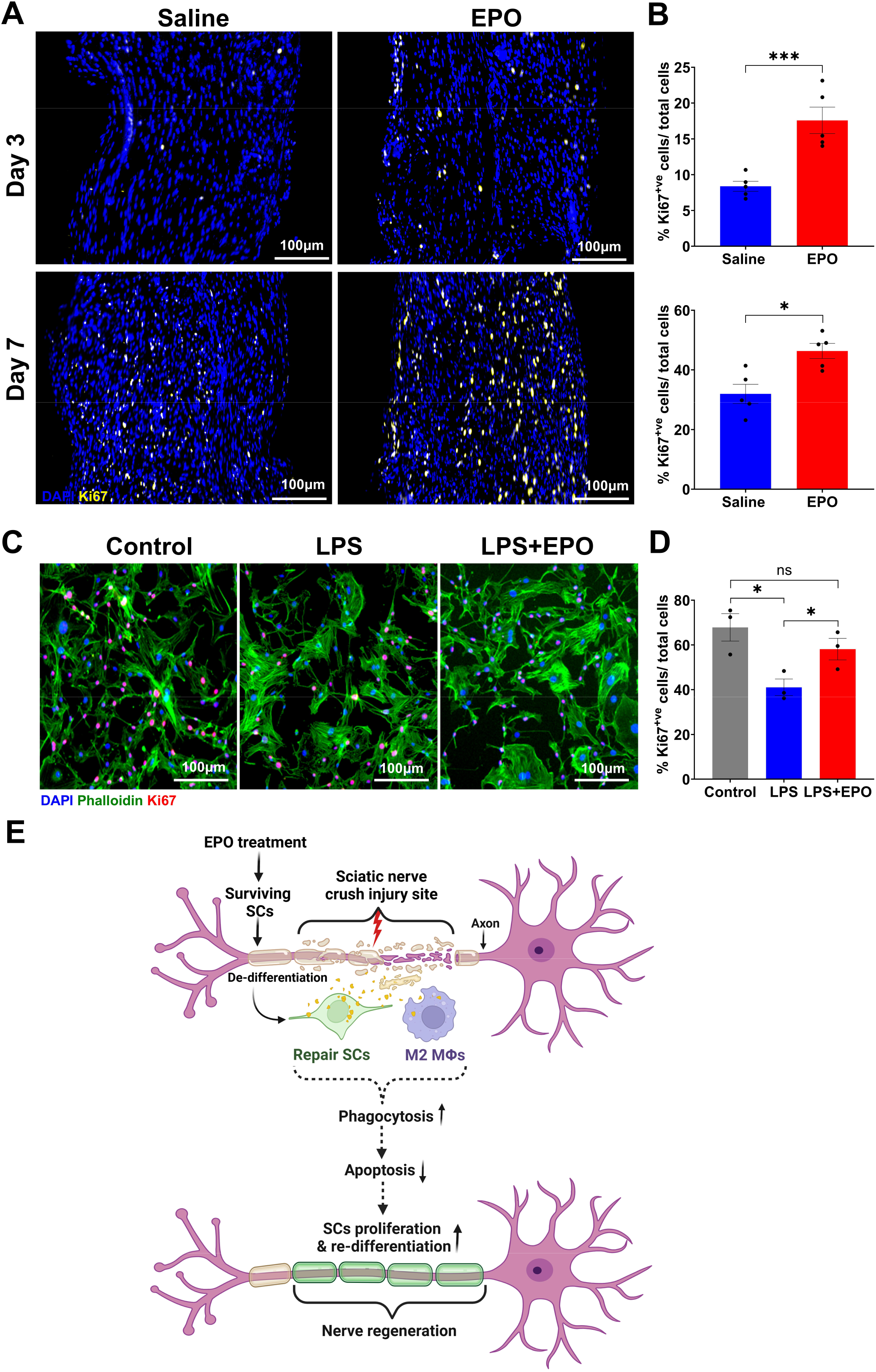
EPO increased cell proliferation in nerve tissues following SNCI. **A, B** Representative IHC images and quantitative cell proliferation results (anti-Ki67 staining) in saline and EPO-treated nerve tissues on post-SNCI days 3, 5, and 7. n = 5/ group. **C, D** Representative IF images and quantitative results of repair SCs (phalloidin staining) % proliferation (Ki67 positive cells/total cells) following 24h EPO (10IU/ mL) treatment under LPS (500ng/ mL) stress conditions. n = 3/ group. **E** A schematic illustration of the role of EPO in rSCs, M2 MΦ myelin phagocytosis, and nerve regeneration following SNCI. Data are represented as mean□±□SEM. The statistical significance is indicated by asterisks (*P□<□0.05 and ***P < 0.0002 vs. saline group) and compared using two-tailed, unpaired t-tests or ordinary one-way ANOVA.

We also investigated EPO’s role in SC proliferation using an *in-vitro* cell culture study under LPS stress conditions by IF staining (Fig. 8C). Our data confirmed that EPO significantly enhanced the proliferation of SCs (Ki67-positive cells) under LPS stress conditions when compared to LPS treatment alone (58.11 ± 4.83 vs. 41.04 ± 3.71; Fig. 8C, D; *P < 0.05). Our findings, from both *in-vivo* and *in-vitro* studies demonstrate a role for EPO in the proliferation of SCs, which may enhance phagocytosis that supports the result found with nerve tissue bulk RNA sequencing. A schematic illustration of the role of EPO in rSCs, M2 MΦ myelin phagocytosis, and nerve regeneration following SNCI is shown in Fig. 8E.

## DISCUSSION

Peripheral nerve injuries are common and yet effective treatments for nerve trauma remain elusive^35^. SCs play a crucial role in repairing injured nerves following PNCI^6,36,37^. During this process, surviving resident myelinating SCs undergo significant molecular changes and differentiate into rSCs^38,39^. These rSCs clear myelin and cellular debris, promoting cell survival, and facilitating axonal regeneration^40,41^. The formation of rSCs after PNCI involves the downregulation of myelination genes and upregulation of repair genes such as c-Jun and p75-NTR^42^. Once activated, rSCs break down redundant or damaged myelin sheaths by initiating proteolysis and myelin autophagy, while recruiting MΦs to the injury site^41,43^.

Our previous work demonstrated that EPO enhanced the transition of M1 to M2 MΦs, as well as the anti-apoptotic effects and phagocytosis of myelin debris after SNCI^1^. However, the significance of EPO on rSC debris clearance remained unknown, despite some encouraging clinical translation of our findings and the findings of others. Recent work also highlights the importance of both rSCs and MΦs in cellular debris phagocytosis and nerve regeneration^15,21,24^. However, few studies specifically investigated SNCIs using mice^15,44^, and none assessed the benefits of EPO under these conditions using transcriptome analysis. We hypothesized that EPO may affect the activity of SCs and MΦs in the clearance of myelin, while also reducing apoptosis through effective trans-differentiation at the injury site following SNCI.

In the current study, we used a calibrated SNCI mouse model^27^ to examine cellular and molecular changes at the injury site through bulk RNA sequencing at early (day 3) and intermediate (days 5 and 7) post-injury time points. Our data analysis indicated that EPO significantly modulated SC and MΦ trans-differentiation, apoptosis, phagocytosis, autophagy, and neuro-regenerative biological pathways following SNCI. We confirmed that EPO protects against apoptosis by enhancing the transformation of SCs and MΦs, which accelerates myelin debris phagocytosis at the nerve injury site after SNCI. *In-vitro*, EPO accelerates the transition of SCs to rSCs, promotes SC proliferation, and enhances phagocytosis of myelin and dead SCs under LPS stress.

We understand that after SNCI, resident SCs are the first to initiate myelin fragmentation, which leads to the breakdown of large myelin fragments and enhances rSC phagocytosis along with infiltrating MΦ phagocytosis. Our bulk RNA sequencing analyses revealed that EPO significantly upregulated genes associated with pathways that promote lysosomal or proteasomal protein catabolism and autophagy, as well as endocytosis and opsonization, on post-SNCI days 3, 5, and 7. Interestingly, the expression levels of specific genes associated with these pathways varied at each time point. We and others have shown that on day 3 post-SNCI, large intact myelin fragments were present at the injury site, along with broken myelin that initiated myelin phagocytosis by rSCs (p75-NTR positive cells). By day 5 post-SNCI, rSCs significantly cleared debris, whereas phagocytic activity decreased on day 7. This reduction was attributed to the efficient early clearance of debris influenced by EPO treatment. We also showed EPO has a significant role in accelerating M2 (CD206 positive) MΦ myelin phagocytosis on day 5.

Several studies support the notion that the early clearance of myelin debris enhances nerve regeneration by reducing apoptosis^45–47^. Our DAB-TUNEL data showed that EPO treatment significantly reduced apoptosis. This finding was further supported by our bulk RNA sequencing analyses conducted on post-SNCI days 3, 5, and 7, which revealed a significant increase in apoptosis in the saline-treated group. Both rSCs and MΦs play a crucial role in myelin clearance on day 5 post-SNCI. Together, these studies suggest that day 5 post-SNCI is the peak period for myelin breakdown and phagocytosis, which supports the efficacy of EPO in reducing apoptosis and promoting functional recovery. *In-vitro* SCs results support our *in-vivo* findings that EPO accelerates rSCs phagocytosis and cell differentiation under LPS stress. This aligns with our earlier work on MΦs^1^ and highlights the crucial roles of EPO in augmenting SCs and MΦs activities for nerve regeneration.

EPO’s role in PNCI involves crush site debris clearance. Our previous work found large effects on MΦs^1^, and now we show changes in SC function that is complementary. Bulk RNA transcriptomic data provides a valuable resource for peripheral nerve researchers, particularly for understanding the pathways related to the trans-differentiation of SCs and MΦs in PNCI. EPO significantly enhances phagocytosis of regenerative rSCs and M2 MΦs while reducing apoptosis. Perhaps these distinct roles on debris clearance cells are the reason for EPO mediated effects on PNCI recovery. Further investigation into the individual cellular effects on SCs and MΦs cell types using single-cell RNA sequencing techniques may offer special resolution in the injury site itself to inform our understanding.

## MATERIALS AND METHODS

### Vertebrate animals

Ten-week-old C57BL/6J male mice weighing 25 ± 3 g were procured from Jackson Laboratories (Bar Harbor, ME). All animal experiments were approved by the Institutional Animal Care and Use Committee (IACUC) at The University of Arizona College of Medicine, Tucson, AZ, and Penn State University, Hershey, PA.

### Sciatic nerve crush injury mouse model

Mice were anesthetized by intraperitoneal injection of ketamine hydrochloride (100 mg/kg) and xylazine (10 mg/kg), purchased from Dechra Veterinary Products, KS, USA. The animal hair was removed from the lower lumbar region using a trimmer, and skin was prepped for nerve crush injury using a 70 % alcohol swab (# 5110, Covidien) and 5 % povidone-iodine applications (# NDC67618-155-16, Betadine). The sciatic nerve crush injury (SNCI) was performed using our established method^27^, which is precise and reproducible. After the SNCI, all animals received extended-release buprenorphine (3.25 mg/kg, # NDC86084-100-30, Ethiqa XR, Fidelis Animal Health) subcutaneously as analgesia. The experimental animals were randomly assigned to either whole transcriptome (n = 3 or 4 animals/group) or validation (n = 5 animals/group) studies (groups: normal saline, 0.1 ml/mouse; EPO, 5000 IU/kg b. wt., # NDC 0069-1305-10, Retacrit). EPO or saline was administered intraperitoneally immediately after surgery, then again on post-surgery days 1 and 2. All animals were euthanized using an isoflurane anesthesia followed by cervical dislocation on days 3, 5, and 7. The injured area of the SN was harvested for transcriptome analysis and validation of apoptosis, myelin phagocytosis, SC proliferation, and differentiation using immunohistochemistry (IHC) and Immunofluorescence (IF) staining.

### Nerve tissue harvesting, RNA extraction, library preparation, and RNA sequencing

Injured SN tissues were surgically harvested and rapidly pulverized using a steel plate in liquid nitrogen, then immediately collected into TRIzol reagent (# 15596026, Invitrogen) and stored at -80°C. Total RNA was extracted using a PureLink^TM^ RNAmicro kit (# 12183016, Thermo Fisher Scientific). RNA quantity and quality control (QC) were assessed using an RNA 6000 Pico assay kit (Agilent 2100 Bioanalyzer Systems, CA, USA), and RNA integrity number (RIN) values above 7.5 were selected for analyses. Sequencing libraries were prepared using sparQ RNA-Seq HMR kit (# 95216-096, Quantabio) with an input of 250 ng of RNA. The quality of the final libraries was assessed on a TapeStation 4150 system (Agilent) and quantified using the Qubit DNA HS reagent (# Q32854, Thermo Fisher Scientific). All libraries were diluted to a final concentration of 2 nmol/L and combined into an equimolar pooled library before sequencing. The pooled library was diluted, denatured, and sequenced on an Illumina NovaSeq^TM^ 6000 using an SP flow cell. Fastq files were generated using a Base Space application (BCL Convert v 2.1.0). All the fastq files were imported into the Partek Flow Software Suite (Partek Inc., Chesterfield, MO, USA), where 3’ base trimming was performed to remove reads with Phred quality scores below 20. The trimmed reads were then aligned to the mm39 mouse genome assembly using the Spliced Transcripts Alignment to a Reference (STAR) method^48^ and annotated with the current Ensembl 107 database release. Median ratio normalization was applied, and differential gene expression was analyzed using Bioconductor-DESeq2^49^. Principal Component Analysis (PCA) was conducted for each group to reduce dimensionality and to visualize differences in gene expression profiles that accounted for more than 1 % of the variance in the dataset. Differentially expressed genes (DEGs) were identified through Partek Flow’s gene differential filter analysis, based on a false discovery rate (FDR) of less than 0.05 and a Log2 fold change (FC) greater than 1.5. Functional annotation and gene enrichment analysis were conducted using Partek Flow, which provides pathways obtained from gene set enrichment analysis (GSEA) and ANOVA enrichment. Before performing the GSEA analysis, genes that met the differential gene expression (DGE) thresholds were pre-ranked^50^. The analysis utilized specific gene set collections from the Molecular Signatures Database (MSigDB), including Gene Ontology (GO) Biological Process, KEGG, and Reactome. Significant pathways related to apoptosis, phagocytosis, immune function, and neuro-regeneration were manually curated from the enrichment clusters and assigned to comparisons for days 3, 5, and 7 (untreated injury vs. EPO-treated injury).

### Nerve tissue TUNEL staining□□

Cell death or apoptosis induced by SNCI was assessed using a terminal deoxynucleotidyl transferase dUTP nick end labeling (TUNEL) assay kit (HRP-DAB) (# ab206386, Abcam). The staining procedure followed the methodology outlined in our previous publication^13^. Finally, slides were imaged using a slide scanner (MoticEasyScan, SF, USA) at 80 X magnification. Image analysis and quantification were performed using NIH ImageJ-1.53e software.

### Nerve tissue immunofluorescence staining

Nerve tissue IF staining was performed as described in our previous publication^1^. In brief, antigen retrieval was performed using a 10 mM sodium citrate buffer (pH 6.0) for 20 minutes at 95 °C. Permeabilization and blocking of nonspecific binding were performed using 1 % Triton X-100 and 5 % goat serum, respectively. Next, primary antibody staining was performed using p75-NTR (1:100, # ab1554, Millipore Sigma), MPZ (1:100, # PZO, Aveslabs), Ki67 (1:100, # 9129, Cell Signaling), and c-Jun (1:100, # Sc74543, Santa Cruz) with an overnight incubation at 4 °C. The samples were then washed three times with phosphate-buffered saline (PBS) and incubated at room temperature for one hour with the appropriate secondary antibodies: anti-Rabbit-Alexa Fluor 488 (1:1000, # A11034, Invitrogen), anti-Mouse-Alexa Fluor 488 (1:1000, # A32723, Invitrogen), anti-Chicken-Alexa Fluor 647 (1:1000, # A21449, Invitrogen), and anti-Rabbit-Alexa Fluor 647 (1:1000, # A32733, Invitrogen). Staining without primary antibodies was used as a control for nonspecific fluorescence. Nuclei were counter-stained using ProLong^TM^Gold anti-fade reagent with DAPI (# P36935, Thermo Fisher Scientific), and sections were examined under a fluorescent microscope (# DM6000, Leica, IL, USA). Image analysis and quantification were performed using NIH ImageJ-1.53e software.

### Myelin protein extraction and quantification

Mice were euthanized as described in our methods to harvest brain, and blood stains were removed by rinsing with ice-cold phosphate-buffered saline (1X PBS). The brain tissues were homogenized in 0.32 M sucrose buffer with protease and phosphatase inhibitors (# 78442, Thermo Fisher Scientific) using a handheld motor homogenizer (# Z359971, Sigma) until a smooth consistency^51^. The homogenate was centrifuged at 800 × g for 10 minutes at 4 °C to eliminate nuclei and cell debris. The resulting supernatant was collected and centrifuged at 10,000 × g for 15 minutes at 4 °C to pellet the crude myelin. The myelin pellet was washed in the homogenization buffer and resuspended in 1 mL of PBS. Finally, the protein concentration of the crude myelin was quantified using the bicinchoninic acid (BCA) assay (# 23225, Thermo Fisher Scientific) to conduct SC phagocytosis studies.

### *In-vitro* Schwann cells phagocytosis by immunofluorescence imaging

The phagocytosis of myelin and apoptotic/dead SCs by live SCs with or without LPS (lipopolysaccharides)/ EPO treatment condition was performed using an IF by following our previous method^1^. In brief, the SCs (passage 1) derived from the adult mice SN were seeded into four-well slides (∼3×10^4^ cells per well) with SCs complete medium (# 1701, ScienCell) and incubated in the humidified chamber until they reached 95 % confluence at 37 °C and 5 % CO_2_. SCs were treated with either LPS (500 ng/mL) or LPS (500 ng/mL) + EPO (10 IU/mL) or not treated (control group) for 24 h, and then cells were washed with 1XDPBS and incubated with PKH26 (# MIDI26-1KT, Sigma-Aldrich) labeled myelin (1 mg/ml) or apoptotic SCs (ratio: 1 live SCs:3 dead SCs) for 4h. After incubation, SCs were washed (1XDPBS) and labeled with Flash Phalloidin Green 488 (1:100, # 42420, Biolegend) for intracellular cytoskeleton F-actin staining and coverslips were mounted on glass slides using ProLong^TM^Gold anti-fade reagent with DAPI (# P36935, Thermo Fisher Scientific), for examination under a fluorescent microscope (# DM6000, Leica, IL, USA). The percentages of phagocytosis were calculated using NIH ImageJ-1.53e software by analyzing the ratio of PKH26 to DAPI.

### *In-vitro* Schwann cells phagocytosis by flowcytometry analysis

The phagocytosis study was conducted as described in our previous publication^1^. In brief, SCs (passage 1) were seeded into 60 mm dishes (∼2×10^5^ cells per dish) and incubated in the humidified chamber until they reached 95 % confluence at 37 °C and 5 % CO_2_. LPS +/- EPO treatment (versus healthy untreated SCs) and myelin incubation were performed as described in our publication. Next, single-cell suspensions of SCs were prepared and washed with ice-cold 1X DPBS. Next, cells were resuspended in 1X flow cytometry staining buffer (# FC001, R&D) and were stained with p75-NTR (1:100, # BS-0161R, Bios) conjugated antibody for 30 min. After staining, the cells were resuspended in a staining buffer. The data was acquired using BD FACSDiva™ v7 software (BD FACSCanto II, AZ, USA) and analyzed using FlowJo^TM^ software (Oregon, USA).

### Schwann cells proliferation

SCs were cultured into four-well chamber slides (# 155382PK, Nunc, Lab Tek) at a density of ∼3×10^4^ cells per well with SCs complete medium. The cultured SCs were incubated at 37 °C in 5 % CO_2_ for 24 h. At ∼60 % confluence, the cells were treated with LPS (500 ng/mL) and LPS (500 ng/mL)+EPO (10 IU/mL) for 24 h. Untreated cells served as a control. After treatment, cells were washed with 1XDPBS, fixed with 4 % paraformaldehyde (# J19943.K2, Thermo Fisher Scientific) for 15 min, permeabilized using 0.5 % Triton X-100 for 10 min, and blocked with 5 % BSA for 30 min at room temperature. Next, cells were stained with primary antibody Ki67 (1:200; # 9129, Cell Signaling Technology) and secondary antibody Alexa fluor-647-conjugated Goat anti-rabbit IgG (1:600, # A32733, Thermo Fisher Scientific). Later, cells were stained with Flash Phalloidin Green 488 (1:200, # 4242011, BioLegend) to visualize the F-actin of cells. Finally, coverslips were mounted on glass slides using ProLong^TM^ Diamond antifade mounting medium with DAPI (# P36971, Thermo Fisher Scientific), and cells were observed under a fluorescent microscope (# DM6000, Leica, IL, USA). The percentage of cell proliferation was performed by counting DAPI vs. Ki67 positive cells using NIH ImageJ-1.53e software.

### Protein extraction and Western blotting analysis

Protein extraction and Western blot analyses were performed using our previously published method^52^. In brief, the total protein was extracted from SCs using RIPA lysis buffer (# R0278, Sigma) and quantified using the BCA method (# 23225, Thermo Fisher Scientific). For gel electrophoresis, 50 μg protein was loaded per lane. The proteins were then transferred to a PVDF membrane by wet transfer (# L00686, GenScript). The membrane was blocked with 3 % BSA for 1 h, followed by overnight incubation with primary antibodies (p75-NTR, 1:1000, # AB1554, Sigma; EGR2, 1:3000, # ab245228, Abcam; c-Jun, 1:2000, # 9165S, Cell Signaling Technology; β-actin, 1:5000, # 5125S, Cell Signaling Technology). The secondary antibodies used are an anti-rabbit HRP-linked antibody (# 7074, Cell Signaling Technology) and an anti-mouse HRP-linked antibody (# 7076, Cell Signaling Technology). The blots were developed using Super Signal West Pico PLUS chemiluminescent substrate ECL kit (# 34579, Thermo Fisher Scientific), and images were captured using a G-box ChemiXRQ gel imager. The bands were quantified using NIH ImageJ-1.53e software. All uncut original Western blotting images of targeted proteins are available in the supplemental material.

### Sciatic functional index

To evaluate the sciatic function index (SFI), we performed a walking track analysis (WTA) on post-SNCI days 3, 5, and 7 as described in our previous publication^13^. SFI was calculated using three parameters of footprints: (1) toe spread (TS, first to the fifth toe), (2) total print length (PL), and (3) intermediate toe spread (IT, second to the fourth toe) and the following formula:SFI = −38.3{(EPL-NPL)/NPL} + 109.5{(ETS-NTS)/NTS} + 13.3{(EIT-NIT)/NIT}−8.8, where E for experimental (injured) and N for normal (contralateral uninjured) sides.

### Statistical analysis

All data were analyzed using GraphPad Prism Version 10.1.1 (San Diego, USA). Comparisons between two groups with n ≥ 3 were performed using two-tailed, unpaired t-tests. Ordinary one-way analysis of variance (ANOVA) was used to compare the three groups with n ≥ 3. All values are presented as mean ± SEM. Significance levels (P values < 0.05) were documented using standard symbols (*, **, ***, and **** correspond to P < 0.05, P < 0.0021, P < 0.0002, and P□<□0.0001, respectively).

## ACKNOWLEDGEMENTS

The authors acknowledge The University of Arizona College of Medicine, Tucson, AZ, USA, and The Penn state University, Hershey, PA, USA for supporting this study. We thank Andrew Powell and Dr. Martha Bhattacharya at the University of Arizona Department of Neuroscience for their early contributions to RNA Sequencing analysis and for engaging in discussions during the study.

## AUTHOR CONTRIBUTIONS

PKG (concept and design of the study, animal surgery, data analysis, and interpretation, figure finalization, and manuscript drafting); GE (animal surgery assistance, imaging, tissue collection, processing, and experiments, data acquisition, analysis, and interpretation, and figure generation and finalization); JPH (animal surgery assistance, tissue collection, processing, and experiments, data acquisition, and analysis); AG (data acquisition, analysis, and interpretation); RVG (animal surgery assistance, dosing, tissue collection, processing, and experiments, data acquisition, analysis, and interpretation, and figure generation and finalization); JCE (concept and design of the study, data interpretation, manuscript finalization, and funding acquisition). All authors read and approved the final manuscript.

## ETHICAL APPROVAL

Our manuscript does not contain any human data. Experimental design and animal protocols were approved by the Institutional Animal Care and Use Committee (IACUC) at The University of Arizona College of Medicine, Tucson, AZ, and The Penn State University, Hershey, PA, USA. All experiments were conducted following the approved guidelines and regulations.

## DATA AVAILABILITY

The data presented in this study are available on request from the corresponding author.

