## Supplemental Material for "Erythropoietin decreases apoptosis and promotes Schwann cell repair and phagocytosis following nerve crush injury in mice"

**Prem Kumar Govindappa**

**John C. Elfar**

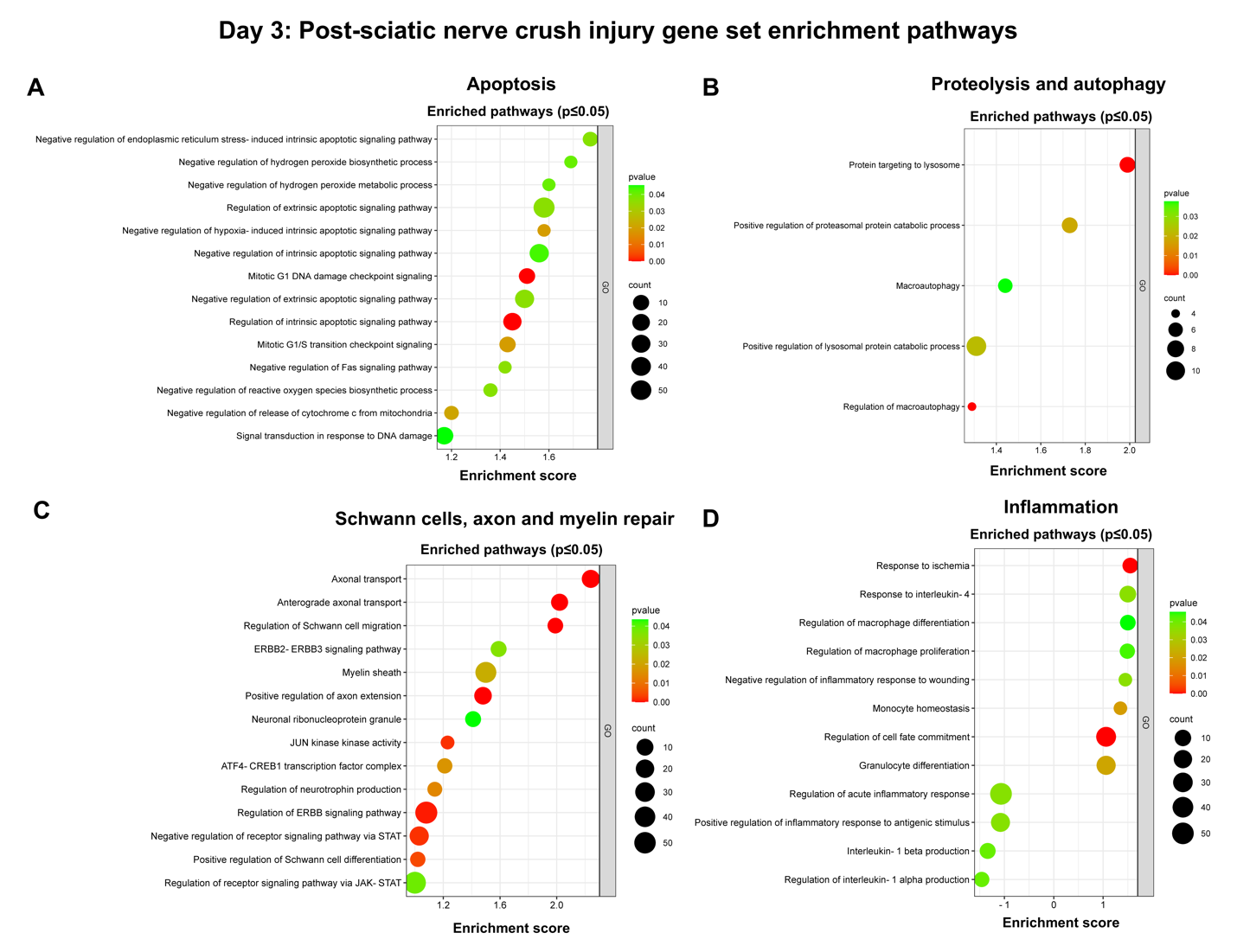
**Supplementary Fig. 1: On Day 3, bulk RNA sequencing revealed EPO-enriched genes for biological pathways in nerves following SNCI. A-D** Gene set enrichment assay (GSEA) pathways are represented on the y-axis with their associated gene numbers (p ≤ 0.05), while the x-axis displays the enrichment score for each pathway. Saline vs. EPO treatment, n = 3/ group.

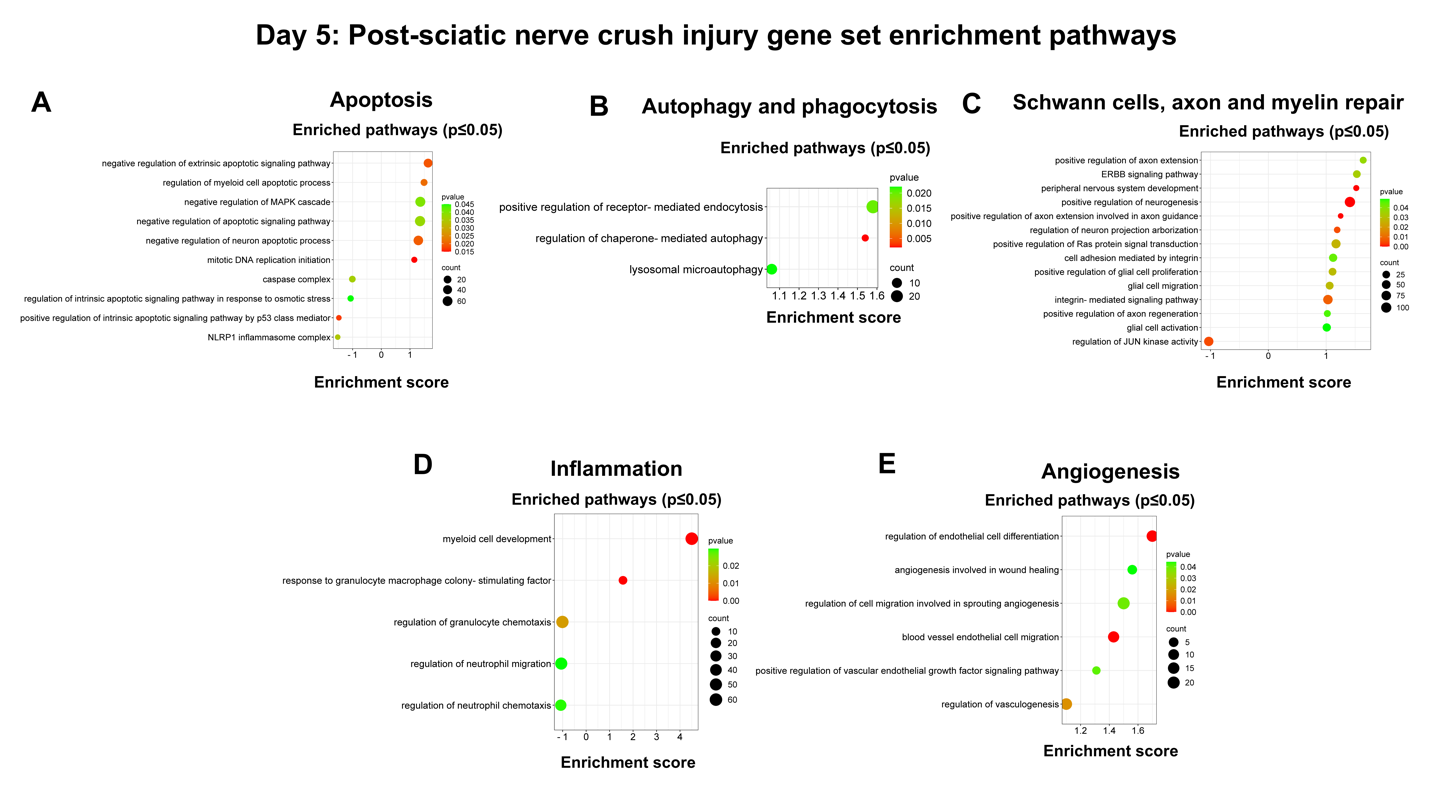
**Supplementary Fig. 2: On Day 5, bulk RNA sequencing revealed EPO-enriched genes for biological pathways in nerves following SNCI. A-E** Gene set enrichment assay (GSEA) pathways are represented on the y-axis with their associated gene numbers (p ≤ 0.05), while the x-axis displays the enrichment score for each pathway. Saline vs. EPO treatment, n = 4/ group.

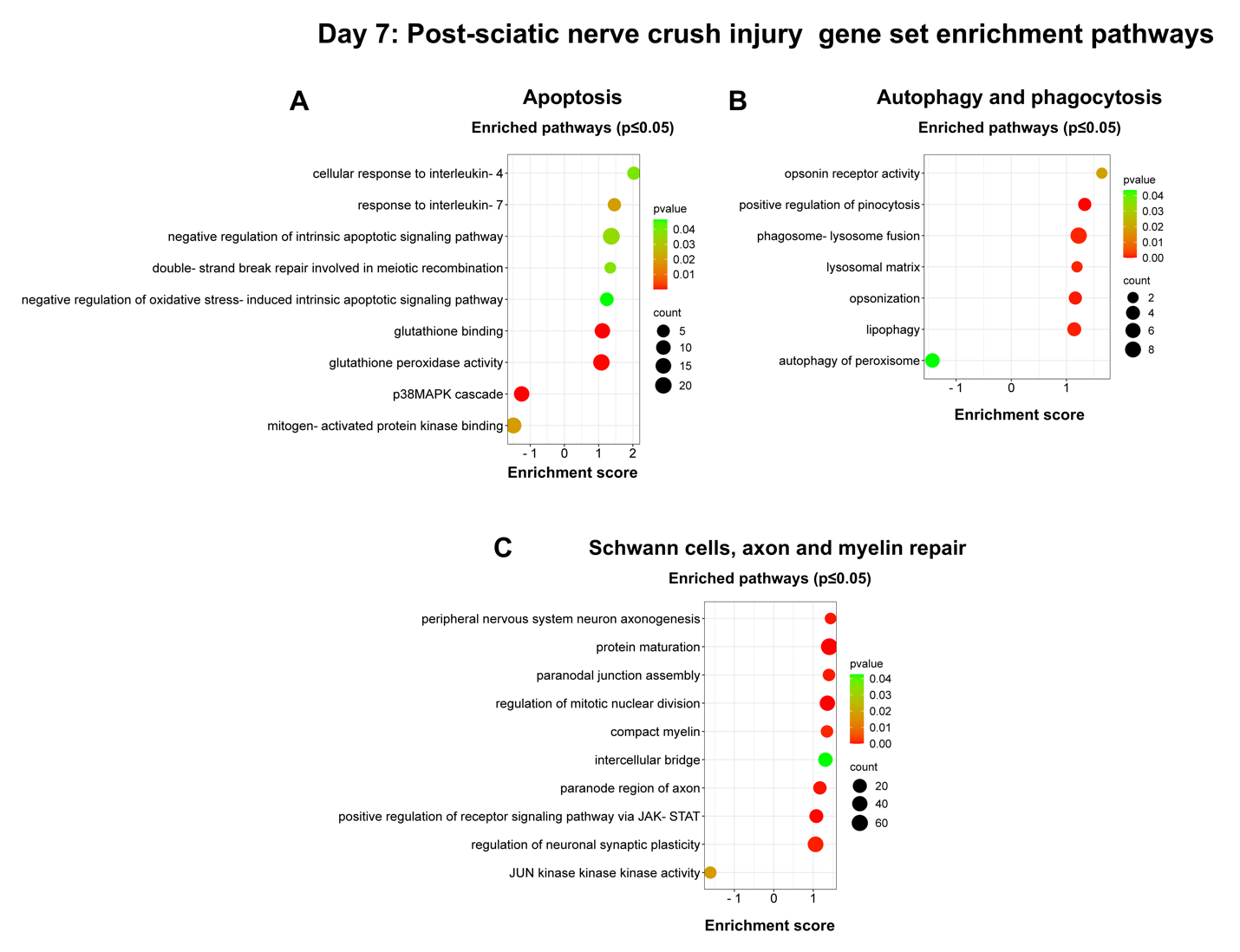
**Supplementary Fig. 3: On Day 7, bulk RNA sequencing revealed EPO-enriched genes for biological pathways in nerves following SNCI. A-E** Gene set enrichment assay (GSEA) pathways are represented on the y-axis with their associated gene numbers (p ≤ 0.05), while the x-axis displays the enrichment score for each pathway. Saline vs. EPO treatment, n = 4/ group.

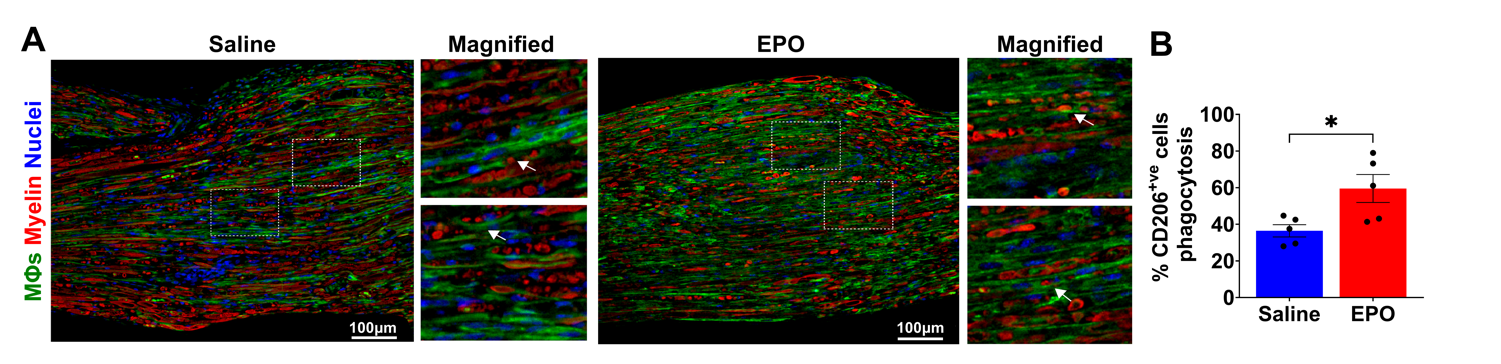
**Supplementary Fig. 4: A, B** Representative IHC images and quantitative results of M2 macrophages (anti-CD206 staining) phagocytosis of myelin debris (anti-MPZ staining) in saline and EPO treated nerve tissues on post-SNCI day 5. n = 5/ group. Data are represented as mean ± SEM. The statistical significance is indicated by asterisks (*P < 0.05 vs. saline group) and compared using two-tailed, unpaired t-tests.

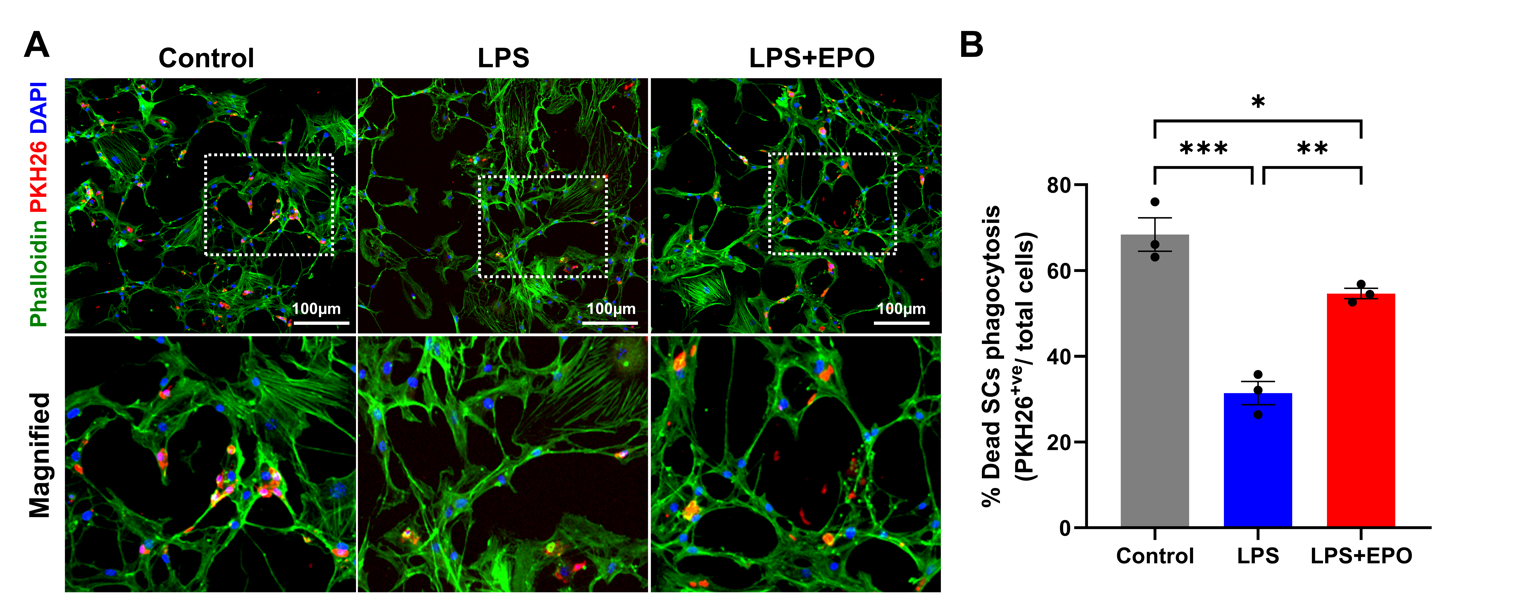
**Supplementary Fig. 5: A, B** Representative IF images and quantitative results of repair SCs (phalloidin staining) phagocytosis of myelin debris (PKH26 staining) following 24h EPO (10IU/ mL) treatment under LPS (500ng/ mL) stress conditions. n = 3/ group. Data are represented as mean ± SEM. The statistical significance is indicated by asterisks (*P < 0.05, **P <  0.0021, and ***P < 0.0002 vs. saline group) and compared using ordinary one-way ANOVA.
