## Supplementary Table 1 for "Erythropoietin decreases apoptosis and promotes Schwann cell repair and phagocytosis following nerve crush injury in mice"

| **Day 3: Saline Vs. EPO (UTI Vs. TI)** | | | | |
| --- | --- | --- | --- | --- |
| **Gene ID** | **Gene name** | **Log2 fold change** | **P-value** | **FDR step up** |
| ENSMUSG00000106438 | Gm32051 | 26.62 | 4.99E-30 | 5.00E-26 |
| ENSMUSG00000091694 | Apol11b | 8.75 | 1.46E-23 | 7.31E-20 |
| ENSMUSG00000073400 | Trim10 | 4.76 | 1.52E-06 | 1.01E-03 |
| ENSMUSG00000027360 | Hdc | 4.02 | 2.71E-06 | 1.43E-03 |
| ENSMUSG00000098650 | Commd1b | 3.03 | 2.24E-07 | 1.87E-04 |
| ENSMUSG00000026580 | Selp | 2.70 | 1.31E-07 | 1.45E-04 |
| ENSMUSG00000027562 | Car2 | 2.60 | 1.99E-08 | 2.85E-05 |
| ENSMUSG00000107092 | Gm7993 | 2.00 | 5.31E-05 | 1.61E-02 |
| ENSMUSG00000061878 | Sphk1 | 1.93 | 9.32E-08 | 1.17E-04 |
| ENSMUSG00000024588 | Fech | 1.90 | 1.57E-07 | 1.47E-04 |
| ENSMUSG00000032860 | P2ry2 | 1.88 | 2.40E-04 | 4.21E-02 |
| ENSMUSG00000025270 | Alas2 | 1.86 | 1.72E-04 | 3.51E-02 |
| ENSMUSG00000022150 | Dab2 | 1.84 | 2.64E-06 | 1.43E-03 |
| ENSMUSG00000030748 | Il4ra | 1.84 | 5.18E-07 | 3.70E-04 |
| ENSMUSG00000069919 | Hba-a1 | 1.82 | 1.17E-04 | 2.60E-02 |
| ENSMUSG00000002289 | Angptl4 | 1.77 | 1.47E-08 | 2.45E-05 |
| ENSMUSG00000069917 | Hba-a2 | 1.77 | 2.61E-04 | 4.51E-02 |
| ENSMUSG00000038729 | Pakap | 1.70 | 2.48E-06 | 1.43E-03 |
| ENSMUSG00000011179 | Odc1 | 1.68 | 1.61E-06 | 1.01E-03 |
| ENSMUSG00000017737 | Mmp9 | 1.68 | 2.94E-04 | 4.96E-02 |
| ENSMUSG00000040809 | Chil3 | 1.65 | 2.17E-05 | 8.05E-03 |
| ENSMUSG00000023272 | Creld2 | 1.63 | 1.03E-04 | 2.51E-02 |
| ENSMUSG00000017009 | Sdc4 | 1.61 | 2.99E-07 | 2.30E-04 |
| ENSMUSG00000063889 | Crem | 1.61 | 2.97E-04 | 4.96E-02 |
| ENSMUSG00000026712 | Mrc1 | 1.59 | 2.30E-04 | 4.21E-02 |
| ENSMUSG00000033885 | Pxk | 1.57 | 2.10E-04 | 3.97E-02 |
| ENSMUSG00000026927 | Entr1 | 1.57 | 6.72E-05 | 1.87E-02 |
| ENSMUSG00000037820 | Tgm2 | 1.54 | 8.46E-05 | 2.23E-02 |
| ENSMUSG00000001403 | Ube2c | 1.50 | 5.85E-05 | 1.72E-02 |
| ENSMUSG00000020484 | Xbp1 | 1.44 | 9.58E-05 | 2.40E-02 |
| ENSMUSG00000008348 | Ubc | 1.42 | 1.13E-04 | 2.57E-02 |
| ENSMUSG00000026970 | Rbms1 | 1.41 | 1.51E-04 | 3.21E-02 |
| ENSMUSG00000118841 | Rn7s2 | -1.41 | 3.55E-05 | 1.19E-02 |
| ENSMUSG00000015355 | Cd48 | -1.45 | 1.77E-04 | 3.52E-02 |
| ENSMUSG00000038543 | BC028528 | -1.47 | 1.83E-04 | 3.52E-02 |
| ENSMUSG00000038811 | Gngt2 | -1.47 | 1.21E-04 | 2.64E-02 |
| ENSMUSG00000026043 | Col3a1 | -1.48 | 6.44E-05 | 1.84E-02 |
| ENSMUSG00000119584 | Rn18s-rs5 | -1.56 | 1.69E-04 | 3.51E-02 |
| ENSMUSG00000024781 | Lipa | -1.62 | 4.76E-05 | 1.49E-02 |
| ENSMUSG00000015568 | Lpl | -1.63 | 1.13E-04 | 2.57E-02 |
| ENSMUSG00000027750 | Postn | -1.66 | 1.08E-04 | 2.57E-02 |
| ENSMUSG00000020053 | Igf1 | -1.73 | 1.90E-05 | 7.31E-03 |
| ENSMUSG00000022665 | Ccdc80 | -1.75 | 3.69E-06 | 1.81E-03 |
| ENSMUSG00000078896 | Zfp965 | -1.77 | 2.37E-04 | 4.21E-02 |
| ENSMUSG00000036594 | H2-Aa | -1.78 | 9.41E-06 | 3.92E-03 |
| ENSMUSG00000059824 | Dbp | -1.85 | 2.32E-04 | 4.21E-02 |
| ENSMUSG00000001348 | Acp5 | -1.88 | 2.51E-05 | 8.98E-03 |
| ENSMUSG00000022371 | Col14a1 | -1.89 | 1.61E-07 | 1.47E-04 |
| ENSMUSG00000030147 | Clec4b1 | -1.97 | 1.32E-05 | 5.29E-03 |
| ENSMUSG00000031443 | F7 | -2.05 | 9.33E-05 | 2.39E-02 |
| ENSMUSG00000071042 | Rasgrp3 | -2.08 | 7.15E-05 | 1.93E-02 |
| ENSMUSG00000040913 | Fbxw4 | -2.24 | 1.02E-08 | 2.05E-05 |
| ENSMUSG00000000440 | Pparg | -2.25 | 4.70E-05 | 1.49E-02 |
| ENSMUSG00000033544 | Angptl1 | -2.30 | 3.79E-06 | 1.81E-03 |
| ENSMUSG00000042254 | Cilp | -2.33 | 5.08E-06 | 2.31E-03 |
| ENSMUSG00000016262 | Sertad4 | -2.38 | 8.57E-06 | 3.73E-03 |
| ENSMUSG00000021388 | Aspn | -2.44 | 1.14E-09 | 3.81E-06 |
| ENSMUSG00000029675 | Eln | -2.58 | 1.83E-04 | 3.52E-02 |
| ENSMUSG00000043629 | 1700019D03Rik | -2.61 | 3.48E-05 | 1.19E-02 |
| ENSMUSG00000042436 | Mfap4 | -2.92 | 4.81E-09 | 1.20E-05 |
