## Supplementary Table 2 for "Erythropoietin decreases apoptosis and promotes Schwann cell repair and phagocytosis following nerve crush injury in mice"

| **Day 3: Myeloid and inflammation regulatory genes** | | | |
| --- | --- | --- | --- |
| **Gene ID** | **Gene name** | **Fold change (D3TI vs D3UTI)** | **P-value (D3TI vs D3UTI)** |
| ENSMUSG00000038067 | Csf3 | 11.57 | 0.00 |
| ENSMUSG00000006574 | Slc4a1 | 5.45 | 0.00 |
| ENSMUSG00000005672 | Kit | 2.48 | 0.04 |
| ENSMUSG00000032115 | Hyou1 | 1.91 | 0.01 |
| ENSMUSG00000025270 | Alas2 | 1.86 | 0.00 |
| ENSMUSG00000069919 | Hba-a1 | 1.82 | 0.00 |
| ENSMUSG00000069917 | Hba-a2 | 1.77 | 0.00 |
| ENSMUSG00000036986 | Pml | 1.73 | 0.00 |
| ENSMUSG00000026712 | Mrc1 | 1.59 | 0.00 |
| ENSMUSG00000038871 | Bpgm | 1.55 | 0.01 |
| ENSMUSG00000025473 | Adam8 | 1.50 | 0.00 |
| ENSMUSG00000020484 | Xbp1 | 1.44 | 0.00 |
| ENSMUSG00000002870 | Mcm2 | 1.43 | 0.02 |
| ENSMUSG00000022951 | Rcan1 | 1.43 | 0.00 |
| ENSMUSG00000026864 | Hspa5 | 1.41 | 0.00 |
| ENSMUSG00000007659 | Bcl2l1 | 1.41 | 0.01 |
| ENSMUSG00000024810 | Il33 | 1.39 | 0.01 |
| ENSMUSG00000071646 | Mta2 | 1.39 | 0.03 |
| ENSMUSG00000079227 | Ccr5 | 1.36 | 0.05 |
| ENSMUSG00000112449 | Srp54b | 1.29 | 0.04 |
| ENSMUSG00000015947 | Fcgr1 | 1.29 | 0.02 |
| ENSMUSG00000004040 | Stat3 | 1.29 | 0.01 |
| ENSMUSG00000056749 | Nfil3 | 1.28 | 0.05 |
| ENSMUSG00000024539 | Ptpn2 | 1.27 | 0.03 |
| ENSMUSG00000021686 | Ap3b1 | 1.27 | 0.05 |
| ENSMUSG00000059552 | Trp53 | 1.27 | 0.04 |
| ENSMUSG00000042228 | Lyn | 1.26 | 0.04 |
| ENSMUSG00000023004 | Tuba1b | 1.24 | 0.01 |
| ENSMUSG00000023944 | Hsp90ab1 | 1.18 | 0.05 |
| ENSMUSG00000032786 | Alas1 | -1.38 | 0.01 |
| ENSMUSG00000026548 | Slamf9 | -1.41 | 0.01 |
| ENSMUSG00000030117 | Gdf3 | -1.42 | 0.02 |
| ENSMUSG00000015133 | Lrrk1 | -1.43 | 0.04 |
| ENSMUSG00000019326 | Aoc3 | -1.67 | 0.02 |
| ENSMUSG00000024164 | C3 | -1.88 | 0.00 |
| ENSMUSG00000001348 | Acp5 | -1.88 | 0.00 |
| ENSMUSG00000067201 | H2-M9 | -2.03 | 0.01 |
| ENSMUSG00000000440 | Pparg | -2.25 | 0.00 |
| ENSMUSG00000005339 | Fcer1a | -2.36 | 0.02 |
| ENSMUSG00000027996 | Sfrp2 | -2.54 | 0.05 |
| ENSMUSG00000026012 | Cd28 | -4.61 | 0.00 |
| ENSMUSG00000042429 | Adora1 | -4.82 | 0.01 |

| **Day 3: Apoptosis regulatory genes** | | | |
| --- | --- | --- | --- |
| **Gene ID** | **Gene name** | **Fold change (D3TI vs D3UTI)** | **P-value (D3TI vs D3UTI)** |
| ENSMUSG00000021070 | Bdkrb2 | 2.60 | 0.01 |
| ENSMUSG00000062991 | Nrg1 | 2.09 | 0.04 |
| ENSMUSG00000032115 | Hyou1 | 1.91 | 0.01 |
| ENSMUSG00000022150 | Dab2 | 1.84 | 0.00 |
| ENSMUSG00000026826 | Nr4a2 | 1.74 | 0.03 |
| ENSMUSG00000036986 | Pml | 1.73 | 0.00 |
| ENSMUSG00000017737 | Mmp9 | 1.68 | 0.00 |
| ENSMUSG00000036432 | Siah2 | 1.54 | 0.02 |
| ENSMUSG00000032350 | Gclc | 1.52 | 0.05 |
| ENSMUSG00000036980 | Taf6 | 1.50 | 0.01 |
| ENSMUSG00000020484 | Xbp1 | 1.44 | 0.00 |
| ENSMUSG00000019969 | Psen1 | 1.41 | 0.00 |
| ENSMUSG00000053110 | Yap1 | 1.41 | 0.04 |
| ENSMUSG00000007659 | Bcl2l1 | 1.41 | 0.01 |
| ENSMUSG00000021109 | Hif1a | 1.37 | 0.02 |
| ENSMUSG00000028820 | Sfpq | 1.35 | 0.02 |
| ENSMUSG00000095567 | Noc2l | 1.34 | 0.04 |
| ENSMUSG00000006301 | Tmbim1 | 1.33 | 0.00 |
| ENSMUSG00000031078 | Cttn | 1.33 | 0.01 |
| ENSMUSG00000028063 | Lmna | 1.32 | 0.00 |
| ENSMUSG00000078652 | Psme3 | 1.30 | 0.03 |
| ENSMUSG00000027111 | Itga6 | 1.30 | 0.04 |
| ENSMUSG00000024927 | Rela | 1.29 | 0.04 |
| ENSMUSG00000004936 | Map2k1 | 1.28 | 0.01 |
| ENSMUSG00000024539 | Ptpn2 | 1.27 | 0.03 |
| ENSMUSG00000059552 | Trp53 | 1.27 | 0.04 |
| ENSMUSG00000005312 | Ubqln1 | 1.26 | 0.04 |
| ENSMUSG00000027540 | Ptpn1 | 1.26 | 0.03 |

| **Day 3: Proteolysis and autophagy regulatory genes** | | | |
| --- | --- | --- | --- |
| **Gene ID** | **Gene name** | **Fold change (D3TI vs D3UTI)** | **P-value (D3TI vs D3UTI)** |
| ENSMUSG00000022150 | Dab2 | 1.84 | 0.00 |
| ENSMUSG00000038058 | Nod1 | 1.69 | 0.01 |
| ENSMUSG00000033885 | Pxk | 1.57 | 0.00 |
| ENSMUSG00000032350 | Gclc | 1.52 | 0.05 |
| ENSMUSG00000002845 | Tmem39a | 1.51 | 0.00 |
| ENSMUSG00000042364 | Snx18 | 1.42 | 0.04 |
| ENSMUSG00000019969 | Psen1 | 1.41 | 0.00 |
| ENSMUSG00000024810 | Il33 | 1.39 | 0.01 |
| ENSMUSG00000021109 | Hif1a | 1.37 | 0.02 |
| ENSMUSG00000015656 | Hspa8 | 1.36 | 0.00 |
| ENSMUSG00000037331 | Larp1 | 1.36 | 0.03 |
| ENSMUSG00000030867 | Plk1 | 1.35 | 0.02 |
| ENSMUSG00000030847 | Bag3 | 1.34 | 0.04 |
| ENSMUSG00000025793 | Hgs | 1.34 | 0.04 |
| ENSMUSG00000020235 | Fzr1 | 1.28 | 0.03 |
| ENSMUSG00000003813 | Rad23a | 1.27 | 0.04 |
| ENSMUSG00000021686 | Ap3b1 | 1.27 | 0.05 |
| ENSMUSG00000059552 | Trp53 | 1.27 | 0.04 |
| ENSMUSG00000005312 | Ubqln1 | 1.26 | 0.04 |
| ENSMUSG00000074247 | Dda1 | 1.21 | 0.03 |
| ENSMUSG00000079477 | Rab7 | 1.19 | 0.04 |

| **Day 3: Schwann cells, axon, and myelin repair regulatory genes** | | | |
| --- | --- | --- | --- |
| **Gene ID** | **Gene name** | **Fold change (D3TI vs D3UTI)** | **P-value (D3TI vs D3UTI)** |
| ENSMUSG00000029371 | Cxcl5 | 3.31 | 0.02 |
| ENSMUSG00000027562 | Car2 | 2.60 | 0.00 |
| ENSMUSG00000005672 | Kit | 2.48 | 0.04 |
| ENSMUSG00000046733 | Gprc5a | 2.10 | 0.03 |
| ENSMUSG00000062991 | Nrg1 | 2.09 | 0.04 |
| ENSMUSG00000069919 | Hba-a1 | 1.82 | 0.00 |
| ENSMUSG00000031391 | L1cam | 1.69 | 0.04 |
| ENSMUSG00000035890 | Rnf126 | 1.69 | 0.01 |
| ENSMUSG00000017737 | Mmp9 | 1.68 | 0.00 |
| ENSMUSG00000063889 | Crem | 1.61 | 0.00 |
| ENSMUSG00000032327 | Stra6 | 1.58 | 0.05 |
| ENSMUSG00000027859 | Ngf | 1.53 | 0.04 |
| ENSMUSG00000053175 | Bcl3 | 1.52 | 0.00 |
| ENSMUSG00000024772 | Ehd1 | 1.52 | 0.00 |
| ENSMUSG00000028268 | Gbp3 | 1.50 | 0.04 |
| ENSMUSG00000020476 | Dbnl | 1.46 | 0.00 |
| ENSMUSG00000028034 | Fubp1 | 1.43 | 0.00 |
| ENSMUSG00000008348 | Ubc | 1.42 | 0.00 |
| ENSMUSG00000020027 | Socs2 | 1.42 | 0.01 |
| ENSMUSG00000019969 | Psen1 | 1.41 | 0.00 |
| ENSMUSG00000026864 | Hspa5 | 1.41 | 0.00 |
| ENSMUSG00000032504 | Pdcd6ip | 1.40 | 0.01 |
| ENSMUSG00000053113 | Socs3 | 1.39 | 0.01 |
| ENSMUSG00000034675 | Dbn1 | 1.38 | 0.01 |
| ENSMUSG00000052397 | Ezr | 1.37 | 0.00 |
| ENSMUSG00000021109 | Hif1a | 1.37 | 0.02 |
| ENSMUSG00000024966 | Stip1 | 1.37 | 0.00 |
| ENSMUSG00000015656 | Hspa8 | 1.36 | 0.00 |
| ENSMUSG00000015291 | Gdi1 | 1.36 | 0.01 |
| ENSMUSG00000033352 | Map2k4 | 1.36 | 0.02 |
| ENSMUSG00000051391 | Ywhag | 1.35 | 0.03 |
| ENSMUSG00000033295 | Ptprf | 1.35 | 0.05 |
| ENSMUSG00000025793 | Hgs | 1.34 | 0.04 |
| ENSMUSG00000055407 | Map6 | 1.32 | 0.03 |
| ENSMUSG00000052727 | Map1b | 1.31 | 0.04 |
| ENSMUSG00000017781 | Pitpna | 1.30 | 0.01 |
| ENSMUSG00000062825 | Actg1 | 1.30 | 0.01 |
| ENSMUSG00000028967 | Errfi1 | 1.30 | 0.03 |
| ENSMUSG00000030681 | Mvp | 1.29 | 0.02 |
| ENSMUSG00000032279 | Idh3a | 1.29 | 0.02 |
| ENSMUSG00000025980 | Hspd1 | 1.29 | 0.02 |
| ENSMUSG00000015714 | Cers2 | 1.28 | 0.01 |
| ENSMUSG00000004936 | Map2k1 | 1.28 | 0.01 |
| ENSMUSG00000024539 | Ptpn2 | 1.27 | 0.03 |
| ENSMUSG00000026426 | Arl8a | 1.27 | 0.02 |
| ENSMUSG00000021686 | Ap3b1 | 1.27 | 0.05 |
| ENSMUSG00000005103 | Wdr1 | 1.26 | 0.02 |
| ENSMUSG00000004789 | Dlst | 1.25 | 0.05 |
| ENSMUSG00000023004 | Tuba1b | 1.24 | 0.01 |
| ENSMUSG00000020358 | Hnrnpab | 1.24 | 0.02 |
| ENSMUSG00000022565 | Plec | 1.22 | 0.05 |
| ENSMUSG00000061904 | Slc25a3 | 1.21 | 0.02 |
| ENSMUSG00000036752 | Tubb4b | 1.20 | 0.03 |
| ENSMUSG00000072235 | Tuba1a | 1.20 | 0.04 |
| ENSMUSG00000079477 | Rab7 | 1.19 | 0.04 |
| ENSMUSG00000000440 | Pparg | -2.25 | 0.00 |
| ENSMUSG00000056457 | Prl2c3 | -2.26 | 0.02 |
| ENSMUSG00000079092 | Prl2c2 | -2.62 | 0.01 |
