## Supplementary Table 3 for "Erythropoietin decreases apoptosis and promotes Schwann cell repair and phagocytosis following nerve crush injury in mice"

| **Day 5: Saline Vs. EPO (UTI Vs. TI)** | | | | |
| --- | --- | --- | --- | --- |
| **Gene ID** | **Gene name** | **Log2 fold change** | **P-value** | **FDR step up** |
| ENSMUSG00000091694 | Apol11b | 29.02 | 1.19E-27 | 1.00E-23 |
| ENSMUSG00000023995 | Tspo2 | 16.83 | 1.21E-24 | 5.09E-21 |
| ENSMUSG00000026532 | Spta1 | 15.70 | 6.90E-08 | 3.42E-05 |
| ENSMUSG00000051839 | Gypa | 15.01 | 8.01E-15 | 1.23E-11 |
| ENSMUSG00000083457 | Cyp4b1-ps2 | 11.74 | 1.43E-07 | 6.51E-05 |
| ENSMUSG00000023216 | Epb42 | 10.90 | 6.25E-07 | 2.56E-04 |
| ENSMUSG00000027562 | Car2 | 8.47 | 1.84E-54 | 3.10E-50 |
| ENSMUSG00000073940 | Hbb-bt | 7.08 | 8.95E-12 | 1.08E-08 |
| ENSMUSG00000025270 | Alas2 | 6.91 | 2.30E-08 | 1.44E-05 |
| ENSMUSG00000073063 | Hbq1b | 6.62 | 1.91E-16 | 3.56E-13 |
| ENSMUSG00000025889 | Snca | 6.00 | 4.02E-17 | 9.68E-14 |
| ENSMUSG00000106438 | Gm32051 | 5.94 | 2.67E-08 | 1.60E-05 |
| ENSMUSG00000075184 | F930017D23Rik | 5.79 | 1.63E-06 | 6.55E-04 |
| ENSMUSG00000020641 | Rsad2 | 5.18 | 1.37E-08 | 9.59E-06 |
| ENSMUSG00000052305 | Hbb-bs | 5.07 | 6.81E-08 | 3.42E-05 |
| ENSMUSG00000001763 | Tspan33 | 5.00 | 4.89E-05 | 1.31E-02 |
| ENSMUSG00000038871 | Bpgm | 4.95 | 2.59E-10 | 2.56E-07 |
| ENSMUSG00000034248 | Slc25a37 | 4.81 | 1.84E-21 | 6.18E-18 |
| ENSMUSG00000069917 | Hba-a2 | 4.72 | 9.08E-08 | 4.24E-05 |
| ENSMUSG00000069919 | Hba-a1 | 4.66 | 2.12E-07 | 9.22E-05 |
| ENSMUSG00000039236 | Isg20 | 4.57 | 1.04E-25 | 5.81E-22 |
| ENSMUSG00000024827 | Gldc | 4.38 | 1.75E-04 | 3.69E-02 |
| ENSMUSG00000059602 | Syn3 | 4.16 | 1.52E-05 | 5.02E-03 |
| ENSMUSG00000028906 | Epb41 | 3.54 | 2.12E-15 | 3.57E-12 |
| ENSMUSG00000027078 | Ube2l6 | 3.26 | 6.53E-20 | 1.83E-16 |
| ENSMUSG00000020802 | Ube2o | 3.26 | 5.07E-13 | 6.57E-10 |
| ENSMUSG00000024588 | Fech | 2.93 | 5.45E-14 | 7.64E-11 |
| ENSMUSG00000034449 | Dhrs11 | 2.83 | 2.14E-07 | 9.22E-05 |
| ENSMUSG00000118332 | Fam220a | 2.78 | 2.02E-09 | 1.70E-06 |
| ENSMUSG00000044468 | Tent5c | 2.73 | 5.93E-17 | 1.25E-13 |
| ENSMUSG00000115759 | Gm18787 | 2.56 | 7.50E-06 | 2.68E-03 |
| ENSMUSG00000024442 | Dele1 | 2.16 | 1.70E-04 | 3.67E-02 |
| ENSMUSG00000078139 | AK157302 | 2.12 | 4.14E-11 | 4.65E-08 |
| ENSMUSG00000022367 | Has2 | 2.02 | 1.84E-05 | 5.73E-03 |
| ENSMUSG00000021792 | Prxl2a | 2.02 | 3.00E-09 | 2.41E-06 |
| ENSMUSG00000029922 | Mkrn1 | 1.99 | 4.36E-09 | 3.19E-06 |
| ENSMUSG00000055116 | Arntl | 1.99 | 1.91E-04 | 3.88E-02 |
| ENSMUSG00000044792 | Isca1 | 1.89 | 9.16E-10 | 8.57E-07 |
| ENSMUSG00000079557 | Marchf2 | 1.83 | 3.50E-08 | 1.96E-05 |
| ENSMUSG00000022836 | Mylk | 1.81 | 1.96E-04 | 3.93E-02 |
| ENSMUSG00000021908 | Ncoa4-ps | 1.76 | 1.87E-08 | 1.24E-05 |
| ENSMUSG00000021171 | Esyt2 | 1.71 | 5.58E-05 | 1.47E-02 |
| ENSMUSG00000028381 | Ugcg | 1.67 | 8.86E-08 | 4.24E-05 |
| ENSMUSG00000075254 | Heg1 | 1.66 | 7.16E-05 | 1.80E-02 |
| ENSMUSG00000022893 | Adamts1 | 1.63 | 1.25E-04 | 2.93E-02 |
| ENSMUSG00000030878 | Cdr2 | 1.62 | 1.13E-05 | 3.82E-03 |
| ENSMUSG00000018659 | Pnpo | 1.60 | 9.17E-05 | 2.24E-02 |
| ENSMUSG00000027333 | Smox | 1.58 | 2.97E-08 | 1.72E-05 |
| ENSMUSG00000027187 | Cat | 1.57 | 2.77E-05 | 8.18E-03 |
| ENSMUSG00000022051 | Bnip3l | 1.57 | 1.92E-08 | 1.24E-05 |
| ENSMUSG00000047139 | Cd24a | 1.57 | 1.73E-04 | 3.68E-02 |
| ENSMUSG00000068566 | Myadm | 1.57 | 1.57E-04 | 3.44E-02 |
| ENSMUSG00000021838 | Samd4 | 1.56 | 2.90E-05 | 8.42E-03 |
| ENSMUSG00000023572 | Ccndbp1 | 1.56 | 3.43E-05 | 9.63E-03 |
| ENSMUSG00000056234 | Ncoa4 | 1.55 | 4.12E-06 | 1.54E-03 |
| ENSMUSG00000070867 | Trabd2b | 1.53 | 1.21E-04 | 2.88E-02 |
| ENSMUSG00000073198 | Bnip3l-ps | 1.51 | 2.15E-06 | 8.21E-04 |
| ENSMUSG00000026072 | Il1r1 | 1.47 | 2.08E-04 | 4.08E-02 |
| ENSMUSG00000030282 | Cmas | 1.44 | 9.64E-05 | 2.32E-02 |
| ENSMUSG00000106133 | Gm3724 | 1.44 | 1.58E-04 | 3.44E-02 |
| ENSMUSG00000033306 | Lpp | 1.41 | 1.40E-04 | 3.22E-02 |
| ENSMUSG00000031950 | Gabarapl2 | 1.39 | 1.85E-04 | 3.83E-02 |
| ENSMUSG00000024501 | Dpysl3 | 1.37 | 4.87E-05 | 1.31E-02 |
| ENSMUSG00000055065 | Ddx17 | 1.37 | 3.87E-05 | 1.07E-02 |
| ENSMUSG00000100801 | Gm15459 | 1.35 | 9.28E-06 | 3.19E-03 |
| ENSMUSG00000025151 | Maged1 | 1.31 | 3.11E-05 | 8.88E-03 |
| ENSMUSG00000025268 | Maged2 | 1.30 | 2.48E-04 | 4.68E-02 |
| ENSMUSG00000044258 | Ctla2a | -1.32 | 2.08E-04 | 4.08E-02 |
| ENSMUSG00000024661 | Fth1 | -1.33 | 2.50E-04 | 4.68E-02 |
| ENSMUSG00000074211 | Sdhaf1 | -1.34 | 7.70E-05 | 1.91E-02 |
| ENSMUSG00000024781 | Lipa | -1.38 | 1.71E-05 | 5.52E-03 |
| ENSMUSG00000031765 | Mt1 | -1.42 | 8.27E-06 | 2.90E-03 |
| ENSMUSG00000060586 | H2-Eb1 | -1.43 | 1.56E-04 | 3.44E-02 |
| ENSMUSG00000030147 | Clec4b1 | -1.51 | 1.44E-04 | 3.26E-02 |
| ENSMUSG00000073421 | H2-Ab1 | -1.53 | 6.09E-08 | 3.20E-05 |
| ENSMUSG00000079293 | Clec7a | -1.53 | 1.88E-04 | 3.86E-02 |
| ENSMUSG00000024610 | Cd74 | -1.53 | 1.83E-09 | 1.62E-06 |
| ENSMUSG00000036594 | H2-Aa | -1.57 | 2.47E-07 | 1.04E-04 |
| ENSMUSG00000045193 | Cirbp | -1.65 | 2.35E-04 | 4.49E-02 |
| ENSMUSG00000113769 | 5033406O09Rik | -1.66 | 1.89E-05 | 5.79E-03 |
| ENSMUSG00000035448 | Ccr3 | -2.11 | 4.86E-08 | 2.64E-05 |
| ENSMUSG00000062038 | Gm10108 | -2.32 | 3.86E-09 | 2.95E-06 |
| ENSMUSG00000112129 | Pbld1 | -2.66 | 5.96E-05 | 1.54E-02 |
| ENSMUSG00000057816 | Cfap299 | -2.67 | 6.96E-05 | 1.77E-02 |
| ENSMUSG00000000730 | Dnmt3l | -2.85 | 5.36E-06 | 1.96E-03 |
| ENSMUSG00000031780 | Ccl17 | -2.91 | 2.04E-06 | 7.98E-04 |
| ENSMUSG00000113047 | Gm47469 | -3.11 | 2.20E-04 | 4.26E-02 |
| ENSMUSG00000094388 | Gm8783 | -3.61 | 1.95E-05 | 5.87E-03 |
| ENSMUSG00000039546 | Ajap1 | -4.14 | 1.74E-05 | 5.52E-03 |
| ENSMUSG00000053706 | B430305J03Rik | -7.16 | 5.52E-11 | 5.80E-08 |
