## Supplementary Table 4 for "Erythropoietin decreases apoptosis and promotes Schwann cell repair and phagocytosis following nerve crush injury in mice"

| **Day 5: Myeloid and inflammation regulatory genes** | | | |
| --- | --- | --- | --- |
| **Gene ID** | **Gene name** | **Fold change (D5TI vs D5UTI)** | **P-value (D5TI vs D5UTI)** |
| ENSMUSG00000025270 | Alas2 | 6.91 | 0.00 |
| ENSMUSG00000027398 | Il1b | 5.74 | 0.01 |
| ENSMUSG00000038871 | Bpgm | 4.95 | 0.00 |
| ENSMUSG00000069917 | Hba-a2 | 4.72 | 0.00 |
| ENSMUSG00000069919 | Hba-a1 | 4.66 | 0.00 |
| ENSMUSG00000042265 | Trem1 | 4.41 | 0.03 |
| ENSMUSG00000031543 | Ank1 | 3.75 | 0.01 |
| ENSMUSG00000026180 | Cxcr2 | 3.35 | 0.05 |
| ENSMUSG00000028825 | Rhd | 2.90 | 0.00 |
| ENSMUSG00000071005 | Ccl19 | 2.87 | 0.01 |
| ENSMUSG00000050761 | Gp1bb | 2.63 | 0.00 |
| ENSMUSG00000031616 | Ednra | 1.68 | 0.00 |
| ENSMUSG00000027858 | Tspan2 | 1.64 | 0.04 |
| ENSMUSG00000026072 | Il1r1 | 1.47 | 0.00 |
| ENSMUSG00000004043 | Stat5a | 1.40 | 0.02 |
| ENSMUSG00000031328 | Flna | 1.31 | 0.00 |
| ENSMUSG00000031402 | Mpp1 | 1.27 | 0.01 |
| ENSMUSG00000052384 | Nrros | 1.22 | 0.05 |
| ENSMUSG00000018446 | C1qbp | -1.21 | 0.01 |
| ENSMUSG00000027239 | Mdk | -1.29 | 0.01 |
| ENSMUSG00000024610 | Cd74 | -1.53 | 0.00 |
| ENSMUSG00000031162 | Gata1 | -2.30 | 0.02 |

| **Day 5: Apoptosis regulatory genes** | | | |
| --- | --- | --- | --- |
| **Gene ID** | **Gene name** | **Fold change (D5TI vs D5UTI)** | **P-value (D5TI vs D5UTI)** |
| ENSMUSG00000025889 | Snca | 6.00 | 0.00 |
| ENSMUSG00000027398 | Il1b | 5.74 | 0.01 |
| ENSMUSG00000026180 | Cxcr2 | 3.35 | 0.05 |
| ENSMUSG00000032487 | Ptgs2 | 2.32 | 0.02 |
| ENSMUSG00000102918 | Pcdhgc3 | 2.22 | 0.05 |
| ENSMUSG00000027298 | Tyro3 | 1.98 | 0.04 |
| ENSMUSG00000026235 | Epha4 | 1.98 | 0.02 |
| ENSMUSG00000062991 | Nrg1 | 1.86 | 0.00 |
| ENSMUSG00000068122 | Agtr2 | 1.78 | 0.01 |
| ENSMUSG00000014039 | Prdm15 | 1.64 | 0.04 |
| ENSMUSG00000031980 | Agt | 1.61 | 0.04 |
| ENSMUSG00000022707 | Gbe1 | 1.54 | 0.00 |
| ENSMUSG00000039005 | Tlr4 | 1.50 | 0.00 |
| ENSMUSG00000023951 | Vegfa | 1.48 | 0.00 |
| ENSMUSG00000005087 | Cd44 | 1.47 | 0.01 |
| ENSMUSG00000003541 | Ier3 | 1.46 | 0.02 |
| ENSMUSG00000004043 | Stat5a | 1.40 | 0.02 |
| ENSMUSG00000079227 | Ccr5 | 1.39 | 0.01 |
| ENSMUSG00000029657 | Hsph1 | 1.38 | 0.01 |
| ENSMUSG00000002413 | Braf | 1.36 | 0.04 |
| ENSMUSG00000021868 | Ppif | 1.34 | 0.03 |
| ENSMUSG00000022673 | Mcm4 | 1.34 | 0.01 |
| ENSMUSG00000021756 | Il6st | 1.32 | 0.00 |
| ENSMUSG00000037049 | Smpd1 | 1.32 | 0.00 |
| ENSMUSG00000007659 | Bcl2l1 | 1.31 | 0.00 |
| ENSMUSG00000027177 | Hipk3 | 1.31 | 0.04 |
| ENSMUSG00000041028 | Ghitm | 1.30 | 0.00 |
| ENSMUSG00000020053 | Igf1 | 1.30 | 0.01 |
| ENSMUSG00000020044 | Timp3 | 1.26 | 0.00 |
| ENSMUSG00000032253 | Phip | 1.26 | 0.02 |
| ENSMUSG00000078652 | Psme3 | 1.26 | 0.00 |
| ENSMUSG00000044167 | Foxo1 | 1.24 | 0.05 |
| ENSMUSG00000015839 | Nfe2l2 | 1.23 | 0.01 |
| ENSMUSG00000022564 | Grina | 1.22 | 0.02 |
| ENSMUSG00000109511 | Nup62 | 1.21 | 0.04 |
| ENSMUSG00000022150 | Dab2 | 1.20 | 0.05 |
| ENSMUSG00000024991 | Eif3a | 1.18 | 0.01 |
| ENSMUSG00000028410 | Dnaja1 | 1.17 | 0.03 |
| ENSMUSG00000049086 | Bmyc | -1.20 | 0.01 |
| ENSMUSG00000030793 | Pycard | -1.21 | 0.03 |
| ENSMUSG00000061477 | Rps7 | -1.23 | 0.02 |
| ENSMUSG00000017721 | Pigt | -1.24 | 0.03 |
| ENSMUSG00000025888 | Casp1 | -1.25 | 0.03 |
| ENSMUSG00000028914 | Casp9 | -1.38 | 0.04 |

| **Day 5: Autophagy and phagocytosis regulatory genes** | | | |
| --- | --- | --- | --- |
| **Gene ID** | **Gene name** | **Fold change (D5TI vs D5UTI)** | **P-value (D5TI vs D5UTI)** |
| ENSMUSG00000025889 | Snca | 6.00 | 0.00 |
| ENSMUSG00000071005 | Ccl19 | 2.87 | 0.01 |
| ENSMUSG00000027298 | Tyro3 | 1.98 | 0.04 |
| ENSMUSG00000032308 | Ulk3 | 1.79 | 0.01 |
| ENSMUSG00000023951 | Vegfa | 1.48 | 0.00 |
| ENSMUSG00000019986 | Ahi1 | 1.41 | 0.00 |
| ENSMUSG00000070738 | Dgkd | 1.39 | 0.03 |
| ENSMUSG00000020828 | Pld2 | 1.36 | 0.03 |
| ENSMUSG00000022150 | Dab2 | 1.20 | 0.05 |
| ENSMUSG00000028385 | Snx30 | -1.41 | 0.01 |

| **Day 5: Angiogenesis regulatory genes** | | | |
| --- | --- | --- | --- |
| **Gene ID** | **Gene name** | **Fold change (D5TI vs D5UTI)** | **P-value (D5TI vs D5UTI)** |
| ENSMUSG00000027398 | Il1b | 5.74 | 0.01 |
| ENSMUSG00000019789 | Hey2 | 3.42 | 0.01 |
| ENSMUSG00000032487 | Ptgs2 | 2.32 | 0.02 |
| ENSMUSG00000021823 | Vcl | 1.88 | 0.00 |
| ENSMUSG00000046318 | Ccbe1 | 1.87 | 0.03 |
| ENSMUSG00000027314 | Dll4 | 1.77 | 0.03 |
| ENSMUSG00000044317 | Gpr4 | 1.76 | 0.04 |
| ENSMUSG00000069763 | Tmem100 | 1.63 | 0.05 |
| ENSMUSG00000090698 | Apold1 | 1.48 | 0.02 |
| ENSMUSG00000023951 | Vegfa | 1.48 | 0.00 |
| ENSMUSG00000067586 | S1pr3 | 1.43 | 0.00 |
| ENSMUSG00000038545 | Cul7 | 1.38 | 0.01 |
| ENSMUSG00000022475 | Hdac7 | 1.38 | 0.05 |
| ENSMUSG00000033960 | Jcad | 1.35 | 0.02 |
| ENSMUSG00000024087 | Cyp1b1 | 1.34 | 0.04 |
| ENSMUSG00000023034 | Nr4a1 | 1.32 | 0.05 |
| ENSMUSG00000046768 | Rhoj | 1.30 | 0.01 |
| ENSMUSG00000016128 | Stard13 | 1.28 | 0.02 |
| ENSMUSG00000021796 | Bmpr1a | 1.25 | 0.01 |

| **Day 5: Schwann cells, axon, and myelin repair regulatory genes** | | | |
| --- | --- | --- | --- |
| **Gene ID** | **Gene name** | **Fold change (D5TI vs D5UTI)** | **P-value (D5TI vs D5UTI)** |
| ENSMUSG00000025889 | Snca | 6.00 | 0.00 |
| ENSMUSG00000027398 | Il1b | 5.74 | 0.01 |
| ENSMUSG00000036832 | Lpar3 | 3.52 | 0.01 |
| ENSMUSG00000071005 | Ccl19 | 2.87 | 0.01 |
| ENSMUSG00000045281 | Gpr20 | 2.68 | 0.05 |
| ENSMUSG00000026768 | Itga8 | 2.55 | 0.04 |
| ENSMUSG00000036904 | Fzd8 | 2.00 | 0.00 |
| ENSMUSG00000029999 | Tgfa | 1.99 | 0.01 |
| ENSMUSG00000026235 | Epha4 | 1.98 | 0.02 |
| ENSMUSG00000020681 | Ace | 1.87 | 0.03 |
| ENSMUSG00000062991 | Nrg1 | 1.86 | 0.00 |
| ENSMUSG00000044317 | Gpr4 | 1.76 | 0.04 |
| ENSMUSG00000022231 | Sema5a | 1.75 | 0.00 |
| ENSMUSG00000010797 | Wnt2 | 1.71 | 0.00 |
| ENSMUSG00000031980 | Agt | 1.61 | 0.04 |
| ENSMUSG00000049791 | Fzd4 | 1.56 | 0.02 |
| ENSMUSG00000026271 | Gpr35 | 1.52 | 0.05 |
| ENSMUSG00000039005 | Tlr4 | 1.50 | 0.00 |
| ENSMUSG00000023951 | Vegfa | 1.48 | 0.00 |
| ENSMUSG00000036158 | Prickle1 | 1.46 | 0.05 |
| ENSMUSG00000021379 | Id4 | 1.46 | 0.05 |
| ENSMUSG00000029071 | Dvl1 | 1.45 | 0.02 |
| ENSMUSG00000019997 | Ccn2 | 1.41 | 0.01 |
| ENSMUSG00000027692 | Tnik | 1.39 | 0.04 |
| ENSMUSG00000037362 | Ccn3 | 1.38 | 0.03 |
| ENSMUSG00000038545 | Cul7 | 1.38 | 0.01 |
| ENSMUSG00000021614 | Vcan | 1.38 | 0.00 |
| ENSMUSG00000002413 | Braf | 1.36 | 0.04 |
| ENSMUSG00000061731 | Ext1 | 1.34 | 0.01 |
| ENSMUSG00000026043 | Col3a1 | 1.34 | 0.03 |
| ENSMUSG00000039361 | Picalm | 1.34 | 0.01 |
| ENSMUSG00000034675 | Dbn1 | 1.33 | 0.00 |
| ENSMUSG00000021756 | Il6st | 1.32 | 0.00 |
| ENSMUSG00000027177 | Hipk3 | 1.31 | 0.04 |
| ENSMUSG00000031328 | Flna | 1.31 | 0.00 |
| ENSMUSG00000021065 | Fut8 | 1.31 | 0.01 |
| ENSMUSG00000027204 | Fbn1 | 1.31 | 0.03 |
| ENSMUSG00000020053 | Igf1 | 1.30 | 0.01 |
| ENSMUSG00000039585 | Myo9a | 1.30 | 0.03 |
| ENSMUSG00000021895 | Arhgef3 | 1.29 | 0.04 |
| ENSMUSG00000037820 | Tgm2 | 1.28 | 0.01 |
| ENSMUSG00000028649 | Macf1 | 1.25 | 0.03 |
| ENSMUSG00000020122 | Egfr | 1.21 | 0.05 |
| ENSMUSG00000055407 | Map6 | 1.21 | 0.05 |
| ENSMUSG00000028410 | Dnaja1 | 1.17 | 0.03 |
| ENSMUSG00000030744 | Rps3 | -1.17 | 0.04 |
| ENSMUSG00000025888 | Casp1 | -1.25 | 0.03 |
| ENSMUSG00000040389 | Wdr47 | -1.57 | 0.02 |
| ENSMUSG00000015533 | Itga2 | -1.66 | 0.05 |
| ENSMUSG00000050350 | Gpr18 | -1.76 | 0.04 |
| ENSMUSG00000005947 | Itgae | -3.11 | 0.00 |
