## Supplementary Table 5 for "Erythropoietin decreases apoptosis and promotes Schwann cell repair and phagocytosis following nerve crush injury in mice"

| **Day 7: Saline Vs. EPO (UTI Vs. TI)** | | | | |
| --- | --- | --- | --- | --- |
| **Gene ID** | **Gene name** | **Log2 fold change** | **P-value** | **FDR step up** |
| ENSMUSG00000041216 | Clvs1 | 10.24 | 2.13E-04 | 3.02E-02 |
| ENSMUSG00000056162 | Cndp1 | 3.11 | 3.84E-04 | 3.99E-02 |
| ENSMUSG00000055116 | Arntl | 2.82 | 9.65E-05 | 1.90E-02 |
| ENSMUSG00000039323 | Igfbp2 | 2.81 | 8.00E-05 | 1.72E-02 |
| ENSMUSG00000030048 | Gkn3 | 2.61 | 8.20E-05 | 1.73E-02 |
| ENSMUSG00000032278 | Paqr5 | 2.57 | 9.55E-08 | 1.53E-04 |
| ENSMUSG00000028871 | Rspo1 | 2.48 | 5.67E-05 | 1.61E-02 |
| ENSMUSG00000056749 | Nfil3 | 2.37 | 6.97E-05 | 1.72E-02 |
| ENSMUSG00000027004 | Frzb | 2.36 | 1.32E-06 | 1.14E-03 |
| ENSMUSG00000025804 | Ccr1 | 2.13 | 7.80E-06 | 3.64E-03 |
| ENSMUSG00000059277 | R74862 | 2.12 | 2.81E-04 | 3.43E-02 |
| ENSMUSG00000037379 | Spon2 | 2.10 | 1.29E-06 | 1.14E-03 |
| ENSMUSG00000068762 | Gstm6 | 1.92 | 6.53E-05 | 1.69E-02 |
| ENSMUSG00000071037 | Camkmt | 1.89 | 2.26E-04 | 3.11E-02 |
| ENSMUSG00000027875 | Hmgcs2 | 1.86 | 3.45E-04 | 3.83E-02 |
| ENSMUSG00000036169 | Sostdc1 | 1.85 | 5.03E-04 | 4.65E-02 |
| ENSMUSG00000015224 | Cyp2j9 | 1.82 | 1.21E-04 | 2.19E-02 |
| ENSMUSG00000032060 | Cryab | 1.78 | 7.92E-05 | 1.72E-02 |
| ENSMUSG00000049130 | C5ar1 | 1.77 | 1.96E-05 | 8.15E-03 |
| ENSMUSG00000022766 | Serpind1 | 1.76 | 1.04E-04 | 2.00E-02 |
| ENSMUSG00000042436 | Mfap4 | 1.73 | 7.79E-05 | 1.72E-02 |
| ENSMUSG00000062380 | Tubb3 | 1.71 | 2.75E-04 | 3.43E-02 |
| ENSMUSG00000070371 | Prss36 | 1.70 | 3.24E-04 | 3.70E-02 |
| ENSMUSG00000117257 | Gm4948 | 1.70 | 7.69E-09 | 1.72E-05 |
| ENSMUSG00000037447 | Arid5a | 1.69 | 4.70E-04 | 4.47E-02 |
| ENSMUSG00000049404 | Rarres1 | 1.66 | 1.82E-04 | 2.80E-02 |
| ENSMUSG00000031538 | Plat | 1.66 | 2.58E-07 | 2.89E-04 |
| ENSMUSG00000039911 | Spsb1 | 1.66 | 2.78E-04 | 3.43E-02 |
| ENSMUSG00000072235 | Tuba1a | 1.66 | 2.90E-07 | 2.95E-04 |
| ENSMUSG00000023960 | Enpp5 | 1.65 | 4.57E-04 | 4.45E-02 |
| ENSMUSG00000005505 | Kbtbd4 | 1.65 | 7.09E-05 | 1.72E-02 |
| ENSMUSG00000091269 | Gm6682 | 1.64 | 5.06E-05 | 1.56E-02 |
| ENSMUSG00000031708 | Tecr | 1.63 | 3.44E-09 | 9.64E-06 |
| ENSMUSG00000002845 | Tmem39a | 1.62 | 2.35E-04 | 3.17E-02 |
| ENSMUSG00000063011 | Msln | 1.62 | 3.11E-04 | 3.69E-02 |
| ENSMUSG00000025505 | Tmem80 | 1.61 | 2.62E-06 | 1.83E-03 |
| ENSMUSG00000114547 | Gm3226 | 1.61 | 1.33E-07 | 1.86E-04 |
| ENSMUSG00000091639 | Gm3756 | 1.61 | 1.33E-04 | 2.33E-02 |
| ENSMUSG00000013495 | Tmem175 | 1.60 | 4.31E-06 | 2.35E-03 |
| ENSMUSG00000001473 | Tubb6 | 1.60 | 6.20E-05 | 1.69E-02 |
| ENSMUSG00000066058 | Cldn19 | 1.58 | 4.23E-04 | 4.25E-02 |
| ENSMUSG00000035439 | Haus8 | 1.58 | 2.80E-04 | 3.43E-02 |
| ENSMUSG00000090084 | Srpx | 1.58 | 1.86E-04 | 2.81E-02 |
| ENSMUSG00000058672 | Tubb2a | 1.57 | 6.06E-06 | 3.08E-03 |
| ENSMUSG00000025823 | Pdia4 | 1.57 | 1.60E-07 | 1.99E-04 |
| ENSMUSG00000035372 | 1810055G02Rik | 1.56 | 1.37E-04 | 2.36E-02 |
| ENSMUSG00000038059 | Smim3 | 1.56 | 1.07E-04 | 2.00E-02 |
| ENSMUSG00000058625 | Gm17383 | 1.56 | 4.66E-05 | 1.49E-02 |
| ENSMUSG00000026172 | Bcs1l | 1.54 | 4.47E-04 | 4.43E-02 |
| ENSMUSG00000030894 | Tpp1 | 1.53 | 7.50E-06 | 3.64E-03 |
| ENSMUSG00000020432 | Tcn2 | 1.51 | 1.81E-06 | 1.45E-03 |
| ENSMUSG00000028194 | Ddah1 | 1.50 | 2.54E-04 | 3.35E-02 |
| ENSMUSG00000071035 | Gm5499 | 1.49 | 2.05E-04 | 3.02E-02 |
| ENSMUSG00000053931 | Cnn3 | 1.48 | 1.31E-04 | 2.32E-02 |
| ENSMUSG00000063889 | Crem | 1.48 | 3.62E-04 | 3.96E-02 |
| ENSMUSG00000022257 | Laptm4b | 1.47 | 1.89E-04 | 2.82E-02 |
| ENSMUSG00000027248 | Pdia3 | 1.47 | 4.28E-06 | 2.35E-03 |
| ENSMUSG00000023272 | Creld2 | 1.47 | 1.58E-04 | 2.60E-02 |
| ENSMUSG00000026864 | Hspa5 | 1.47 | 9.23E-06 | 4.13E-03 |
| ENSMUSG00000024875 | Yif1a | 1.47 | 7.74E-05 | 1.72E-02 |
| ENSMUSG00000009894 | Snap47 | 1.46 | 7.80E-05 | 1.72E-02 |
| ENSMUSG00000079111 | Kdelr2 | 1.46 | 3.64E-04 | 3.96E-02 |
| ENSMUSG00000046314 | Stxbp6 | 1.45 | 4.25E-04 | 4.25E-02 |
| ENSMUSG00000021811 | Dnajc9 | 1.44 | 2.18E-04 | 3.05E-02 |
| ENSMUSG00000027804 | Ppid | 1.44 | 1.62E-05 | 7.00E-03 |
| ENSMUSG00000019916 | P4ha1 | 1.43 | 5.76E-05 | 1.61E-02 |
| ENSMUSG00000032123 | Dpagt1 | 1.43 | 4.02E-04 | 4.10E-02 |
| ENSMUSG00000020484 | Xbp1 | 1.43 | 3.68E-05 | 1.33E-02 |
| ENSMUSG00000091421 | Gm4202 | 1.42 | 9.34E-05 | 1.88E-02 |
| ENSMUSG00000023067 | Cdkn1a | 1.42 | 2.45E-04 | 3.27E-02 |
| ENSMUSG00000039887 | Alg14 | 1.41 | 3.90E-04 | 4.01E-02 |
| ENSMUSG00000032116 | Stt3a | 1.41 | 3.82E-04 | 3.99E-02 |
| ENSMUSG00000014402 | Tsg101 | 1.40 | 3.13E-04 | 3.69E-02 |
| ENSMUSG00000082809 | Gm14150 | 1.39 | 4.71E-04 | 4.47E-02 |
| ENSMUSG00000022757 | Tfg | 1.39 | 2.09E-04 | 3.02E-02 |
| ENSMUSG00000021501 | Caml | 1.39 | 4.82E-04 | 4.54E-02 |
| ENSMUSG00000032279 | Idh3a | 1.37 | 2.28E-04 | 3.11E-02 |
| ENSMUSG00000031521 | Aga | 1.37 | 4.65E-04 | 4.47E-02 |
| ENSMUSG00000063902 | Gm7964 | 1.37 | 4.98E-04 | 4.65E-02 |
| ENSMUSG00000003814 | Calr | 1.36 | 1.07E-04 | 2.00E-02 |
| ENSMUSG00000018736 | Ndel1 | 1.36 | 1.77E-04 | 2.80E-02 |
| ENSMUSG00000023904 | Hcfc1r1 | 1.36 | 2.89E-04 | 3.48E-02 |
| ENSMUSG00000016637 | Ift27 | 1.34 | 2.11E-04 | 3.02E-02 |
| ENSMUSG00000030062 | Rpn1 | 1.32 | 5.28E-05 | 1.56E-02 |
| ENSMUSG00000018965 | Ywhah | 1.32 | 2.04E-05 | 8.16E-03 |
| ENSMUSG00000021248 | Tmed10 | 1.32 | 2.81E-04 | 3.43E-02 |
| ENSMUSG00000031029 | Eif3f | -1.38 | 7.30E-05 | 1.72E-02 |
| ENSMUSG00000118866 | Rn7s1 | -1.38 | 3.36E-04 | 3.80E-02 |
| ENSMUSG00000022283 | Pabpc1 | -1.42 | 3.82E-04 | 3.99E-02 |
| ENSMUSG00000062098 | Btbd3 | -1.42 | 5.20E-04 | 4.78E-02 |
| ENSMUSG00000027465 | Tbc1d20 | -1.43 | 3.84E-04 | 3.99E-02 |
| ENSMUSG00000020580 | Rock2 | -1.48 | 2.83E-05 | 1.09E-02 |
| ENSMUSG00000064339 | mt-Rnr2 | -1.50 | 4.40E-06 | 2.35E-03 |
| ENSMUSG00000064337 | mt-Rnr1 | -1.50 | 3.11E-06 | 2.05E-03 |
| ENSMUSG00000045817 | Zfp36l2 | -1.51 | 1.78E-04 | 2.80E-02 |
| ENSMUSG00000022095 | Fam160b2 | -1.53 | 3.85E-04 | 3.99E-02 |
| ENSMUSG00000022053 | Ebf2 | -1.54 | 8.78E-05 | 1.82E-02 |
| ENSMUSG00000089764 | Gm16580 | -1.57 | 4.55E-04 | 4.45E-02 |
| ENSMUSG00000047888 | Tnrc6b | -1.64 | 9.41E-05 | 1.88E-02 |
| ENSMUSG00000044857 | Lemd2 | -1.64 | 3.83E-05 | 1.34E-02 |
| ENSMUSG00000034042 | Gpbp1l1 | -1.70 | 3.21E-04 | 3.70E-02 |
| ENSMUSG00000036452 | Arhgap26 | -1.70 | 1.60E-04 | 2.60E-02 |
| ENSMUSG00000034709 | Ppp1r21 | -1.73 | 2.76E-04 | 3.43E-02 |
| ENSMUSG00000068742 | Cry2 | -1.74 | 2.59E-04 | 3.38E-02 |
| ENSMUSG00000018909 | Arrb1 | -1.78 | 1.20E-04 | 2.19E-02 |
| ENSMUSG00000031133 | Arhgef6 | -1.82 | 4.29E-05 | 1.41E-02 |
| ENSMUSG00000040850 | Psme4 | -1.83 | 3.54E-06 | 2.20E-03 |
| ENSMUSG00000044950 | Pwwp2a | -1.90 | 5.26E-05 | 1.56E-02 |
| ENSMUSG00000017548 | Suz12 | -1.95 | 1.54E-04 | 2.58E-02 |
| ENSMUSG00000009633 | G0s2 | -1.96 | 6.64E-05 | 1.69E-02 |
| ENSMUSG00000037410 | Tbc1d2b | -1.97 | 4.04E-05 | 1.37E-02 |
| ENSMUSG00000056413 | Adap1 | -2.02 | 1.80E-04 | 2.80E-02 |
| ENSMUSG00000030306 | Tmtc1 | -2.02 | 6.53E-05 | 1.69E-02 |
| ENSMUSG00000023078 | Cxcl13 | -2.32 | 1.55E-04 | 2.58E-02 |
| ENSMUSG00000032175 | Tyk2 | -2.44 | 2.20E-06 | 1.64E-03 |
| ENSMUSG00000022389 | Tef | -2.55 | 1.07E-11 | 3.98E-08 |
| ENSMUSG00000059824 | Dbp | -2.65 | 4.66E-13 | 5.22E-09 |
| ENSMUSG00000028957 | Per3 | -2.83 | 8.27E-08 | 1.53E-04 |
| ENSMUSG00000040966 | Slc22a2 | -2.90 | 3.46E-04 | 3.83E-02 |
| ENSMUSG00000041594 | Tmtc4 | -2.93 | 3.23E-04 | 3.70E-02 |
| ENSMUSG00000021775 | Nr1d2 | -3.16 | 9.43E-13 | 5.28E-09 |
| ENSMUSG00000018459 | Slc13a3 | -94.42 | 2.92E-05 | 1.09E-02 |
