## Supplementary Table 6 for "Erythropoietin decreases apoptosis and promotes Schwann cell repair and phagocytosis following nerve crush injury in mice"

| **Day 7: Apoptosis regulatory genes** | | | |
| --- | --- | --- | --- |
| **Gene ID** | **Gene name** | **Fold change (TI vs UTI)** | **P-value (TI vs UTI)** |
| ENSMUSG00000056749 | Nfil3 | 2.37 | 0.00 |
| ENSMUSG00000023826 | Prkn | 1.97 | 0.03 |
| ENSMUSG00000068762 | Gstm6 | 1.92 | 0.00 |
| ENSMUSG00000028690 | Mmachc | 1.64 | 0.00 |
| ENSMUSG00000027323 | Rad51 | 1.64 | 0.00 |
| ENSMUSG00000020377 | Ltc4s | 1.52 | 0.01 |
| ENSMUSG00000027248 | Pdia3 | 1.47 | 0.00 |
| ENSMUSG00000026864 | Hspa5 | 1.47 | 0.00 |
| ENSMUSG00000020484 | Xbp1 | 1.43 | 0.00 |
| ENSMUSG00000096472 | Cdkn2d | 1.40 | 0.00 |
| ENSMUSG00000022037 | Clu | 1.39 | 0.03 |
| ENSMUSG00000028567 | Txndc12 | 1.37 | 0.02 |
| ENSMUSG00000009291 | Pttg1ip | 1.37 | 0.00 |
| ENSMUSG00000059040 | Eno1b | 1.35 | 0.00 |
| ENSMUSG00000060803 | Gstp1 | 1.34 | 0.00 |
| ENSMUSG00000033307 | Mif | 1.33 | 0.00 |
| ENSMUSG00000023004 | Tuba1b | 1.30 | 0.00 |
| ENSMUSG00000063524 | Eno1 | 1.30 | 0.01 |
| ENSMUSG00000103653 | Gstp-ps | 1.30 | 0.01 |
| ENSMUSG00000015839 | Nfe2l2 | 1.30 | 0.00 |
| ENSMUSG00000028597 | Gpx7 | 1.28 | 0.01 |
| ENSMUSG00000058135 | Gstm1 | 1.28 | 0.05 |
| ENSMUSG00000028410 | Dnaja1 | 1.28 | 0.00 |
| ENSMUSG00000006728 | Cdk4 | 1.26 | 0.01 |
| ENSMUSG00000029864 | Gstk1 | 1.24 | 0.04 |
| ENSMUSG00000021771 | Vdac2 | 1.24 | 0.04 |
| ENSMUSG00000024966 | Stip1 | 1.22 | 0.02 |
| ENSMUSG00000026701 | Prdx6 | 1.22 | 0.03 |
| ENSMUSG00000025980 | Hspd1 | 1.21 | 0.04 |
| ENSMUSG00000049792 | Bag5 | 1.20 | 0.04 |
| ENSMUSG00000006005 | Tpr | -1.23 | 0.01 |
| ENSMUSG00000033016 | Nfatc1 | -1.36 | 0.00 |
| ENSMUSG00000005893 | Nr2c2 | -1.45 | 0.05 |
| ENSMUSG00000016528 | Mapkapk2 | -1.48 | 0.01 |
| ENSMUSG00000020063 | Sirt1 | -1.48 | 0.03 |
| ENSMUSG00000030247 | Kcnj8 | -1.50 | 0.02 |
| ENSMUSG00000055024 | Ep300 | -1.70 | 0.00 |
| ENSMUSG00000020400 | Tnip1 | -1.73 | 0.01 |

| **Day 7: Autophagy and phagocytosis regulatory genes** | | | |
| --- | --- | --- | --- |
| **Gene ID** | **Gene name** | **Fold change (TI vs UTI)** | **P-value (TI vs UTI)** |
| ENSMUSG00000037379 | Spon2 | 2.10 | 0.00 |
| ENSMUSG00000049130 | C5ar1 | 1.77 | 0.00 |
| ENSMUSG00000013495 | Tmem175 | 1.60 | 0.00 |
| ENSMUSG00000052688 | Rab7b | 1.27 | 0.02 |
| ENSMUSG00000015656 | Hspa8 | 1.27 | 0.00 |
| ENSMUSG00000028657 | Ppt1 | 1.27 | 0.01 |
| ENSMUSG00000002059 | Rab34 | 1.23 | 0.02 |
| ENSMUSG00000079477 | Rab7 | 1.19 | 0.02 |
| ENSMUSG00000006699 | Cdc42 | 1.19 | 0.04 |
| ENSMUSG00000034218 | Atm | -1.60 | 0.04 |

| **Day 7: Schwann cells, axon, and myelin repair regulatory genes** | | | |
| --- | --- | --- | --- |
| **Gene ID** | **Gene name** | **Fold change (TI vs UTI)** | **P-value (TI vs UTI)** |
| ENSMUSG00000020396 | Nefh | 3.35 | 0.02 |
| ENSMUSG00000022842 | Ece2 | 2.78 | 0.03 |
| ENSMUSG00000026442 | Nfasc | 2.16 | 0.01 |
| ENSMUSG00000021835 | Bmp4 | 2.04 | 0.00 |
| ENSMUSG00000029414 | Kntc1 | 1.95 | 0.04 |
| ENSMUSG00000068762 | Gstm6 | 1.92 | 0.00 |
| ENSMUSG00000062591 | Tubb4a | 1.86 | 0.00 |
| ENSMUSG00000024261 | Syt4 | 1.82 | 0.00 |
| ENSMUSG00000034926 | Dhcr24 | 1.81 | 0.01 |
| ENSMUSG00000062380 | Tubb3 | 1.71 | 0.00 |
| ENSMUSG00000031538 | Plat | 1.66 | 0.00 |
| ENSMUSG00000082361 | Btc | 1.64 | 0.00 |
| ENSMUSG00000001473 | Tubb6 | 1.60 | 0.00 |
| ENSMUSG00000058672 | Tubb2a | 1.57 | 0.00 |
| ENSMUSG00000025823 | Pdia4 | 1.57 | 0.00 |
| ENSMUSG00000045136 | Tubb2b | 1.56 | 0.00 |
| ENSMUSG00000036634 | Mag | 1.53 | 0.02 |
| ENSMUSG00000048574 | Ccnb1-ps | 1.50 | 0.00 |
| ENSMUSG00000039033 | Tasp1 | 1.48 | 0.00 |
| ENSMUSG00000028128 | F3 | 1.47 | 0.01 |
| ENSMUSG00000037031 | Tspan15 | 1.47 | 0.01 |
| ENSMUSG00000037852 | Cpe | 1.45 | 0.01 |
| ENSMUSG00000018217 | Pmp22 | 1.44 | 0.01 |
| ENSMUSG00000017716 | Birc5 | 1.39 | 0.02 |
| ENSMUSG00000041431 | Ccnb1 | 1.39 | 0.01 |
| ENSMUSG00000023000 | Dhh | 1.36 | 0.05 |
| ENSMUSG00000070436 | Serpinh1 | 1.34 | 0.01 |
| ENSMUSG00000022750 | Klhl22 | 1.34 | 0.03 |
| ENSMUSG00000005233 | Spc25 | 1.33 | 0.02 |
| ENSMUSG00000030342 | Cd9 | 1.32 | 0.03 |
| ENSMUSG00000018965 | Ywhah | 1.32 | 0.00 |
| ENSMUSG00000026683 | Nuf2 | 1.31 | 0.02 |
| ENSMUSG00000038943 | Prc1 | 1.30 | 0.02 |
| ENSMUSG00000021190 | Lgmn | 1.29 | 0.02 |
| ENSMUSG00000026029 | Casp8 | 1.29 | 0.01 |
| ENSMUSG00000027978 | Prss12 | 1.29 | 0.05 |
| ENSMUSG00000058135 | Gstm1 | 1.28 | 0.05 |
| ENSMUSG00000024487 | Yipf5 | 1.28 | 0.00 |
| ENSMUSG00000004460 | Dnajb11 | 1.27 | 0.00 |
| ENSMUSG00000020473 | Aebp1 | 1.27 | 0.04 |
| ENSMUSG00000031985 | Gnpat | 1.27 | 0.02 |
| ENSMUSG00000029017 | Pmpcb | 1.26 | 0.01 |
| ENSMUSG00000020737 | Jpt1 | 1.26 | 0.01 |
| ENSMUSG00000037824 | Tspan14 | 1.26 | 0.05 |
| ENSMUSG00000024889 | Rce1 | 1.25 | 0.02 |
| ENSMUSG00000029910 | Mad2l1 | 1.25 | 0.03 |
| ENSMUSG00000015149 | Sirt2 | 1.25 | 0.02 |
| ENSMUSG00000034509 | Mad2l1bp | 1.25 | 0.04 |
| ENSMUSG00000066979 | Bub3 | 1.25 | 0.00 |
| ENSMUSG00000047866 | Lonp2 | 1.25 | 0.03 |
| ENSMUSG00000024537 | Psmg2 | 1.25 | 0.01 |
| ENSMUSG00000078695 | Cisd3 | 1.24 | 0.05 |
| ENSMUSG00000019188 | H13 | 1.23 | 0.01 |
| ENSMUSG00000026889 | Rbm18 | 1.23 | 0.04 |
| ENSMUSG00000043207 | Zmpste24 | 1.23 | 0.05 |
| ENSMUSG00000024847 | Aip | 1.23 | 0.01 |
| ENSMUSG00000001525 | Tubb5 | 1.22 | 0.01 |
| ENSMUSG00000024571 | Naa12 | 1.20 | 0.04 |
| ENSMUSG00000006699 | Cdc42 | 1.19 | 0.04 |
| ENSMUSG00000036752 | Tubb4b | 1.19 | 0.05 |
| ENSMUSG00000032512 | Wdr48 | -1.26 | 0.05 |
| ENSMUSG00000036104 | Rab3gap1 | -1.46 | 0.00 |
| ENSMUSG00000028664 | Ephb2 | -1.47 | 0.05 |
| ENSMUSG00000055024 | Ep300 | -1.70 | 0.00 |
| ENSMUSG00000032089 | Il10ra | -1.83 | 0.00 |
| ENSMUSG00000064128 | Cenpj | -1.93 | 0.02 |
| ENSMUSG00000005672 | Kit | -2.21 | 0.03 |
| ENSMUSG00000032175 | Tyk2 | -2.44 | 0.00 |
| ENSMUSG00000048616 | Nog | -2.95 | 0.02 |
